# Methionine, Not *S*-adenosylmethionine, Acts as a Primary Metabolic Stress Signal for Chromatin Remodeling

**DOI:** 10.64898/2026.05.01.722273

**Authors:** Cassandra M. Leech, Spencer A. Haws, John M. Denu

## Abstract

Epigenetic regulation is tightly linked to cellular metabolism through chromatin-modifying enzymes that depend on central metabolites as co-substrates. Methionine is an essential amino acid that is directly converted by methionine adenosyltransferase 2A (MAT2A) into *S*-adenosylmethionine (SAM), the universal methyl donor required for histone and DNA methylation. Although methionine restriction/depletion can alter the chromatin methylation landscape and improve physiological outcomes in diverse biological systems, it remains unclear whether these effects arise from loss of methionine itself or from secondary depletion of SAM. Here, we show that methionine depletion induces nuclear accumulation of MAT2A together with redistribution of H3K9 methylation, derepression of transposable elements, activation of stress-response pathways, and broad transcriptional reprogramming. Surprisingly, pharmacologic inhibition reduced intracellular SAM to levels comparable to methionine depletion but failed to reproduce these major epigenetic or transcriptional responses. Furthermore, depletion of the SAM-sensor SAMTOR and inhibition of KDM4 histone demethylases did not prevent methionine-dependent chromatin remodeling, indicating that canonical SAM-sensing pathways are not required for this adaptation. Instead, methionine depletion uniquely induced innate immune and integrated stress-response programs consistent with a viral mimicry-like state. These findings demonstrate that methionine availability, rather than SAM abundance, functions as a primary metabolic signal regulating epigenetic adaptation to nutrient stress. Our data support a model in which methionine is sensed independently of SAM abundance and acts upstream of stress signaling pathways that secondarily remodel chromatin.

## Introduction

Chromatin organization, built from repeating units of nucleosomal DNA wrapped around histone octamers, regulates genome accessibility and gene expression. Chemical post-translational modifications (PTMs) placed on histone tails can modulate chromatin structure either directly or indirectly to control transcriptional activity. Chromatin exists along a spectrum from euchromatin, associated with active transcription, to heterochromatin, which is transcriptionally repressive and essential for maintaining genome stability^1,2^. Chromatin states can be highly dynamic, allowing for rapid regulation of gene expression required for normal development, cell-type specificity, stress adaptation, and healthy aging^3–7^.

Heterochromatin comprises a substantial portion (∼45%) of the human genome and is enriched in repetitive elements, including transposable elements (TEs) and satellite DNA. Repetitive transcripts, when expressed, are often deleterious to the cell due to their propensity for forming DNA:RNA hybrids, double-stranded DNA (dsDNA) breaks, or propagating into new genomic regions^8,9^. Long Interspersed Element 1 (LINE1) is one of the most abundant TEs and accounts for nearly 17% of the assembled human genome, though most coding regions have been inactivated by mutations acquired throughout evolution^9–11^. Human Endogenous Retroviruses (HERVs) are another remnant of ancient retroviral infection incorporated predominantly at telomeric and peri-centromeric regions of chromosomes and account for ∼8% of the human genome^12,13^. Proper repression of constitutive heterochromatin, mediated largely by H3K9me3 and DNA methylation, is therefore essential for safeguarding the genome via suppression of satellite and repetitive DNA, including TEs^14–17^.

Epigenetic regulation of chromatin function is closely linked to cellular metabolism, as chromatin-modifying enzymes require metabolite cofactors and co-substrates. Fluctuations in the availability of essential metabolic co-substrates, through changes in diet or disease state, can directly affect a cell’s ability to add or remove these epigenetic modifications^18–21^. Histone PTMs, therefore, function as an integral node connecting metabolism and gene regulation by allowing cells to sense the availability of key metabolites and alter gene expression in real time.

Methionine (Met), is an essential dietary amino acid required for major cellular processes including protein synthesis, redox regulation, RNA processing, post-translation modifications, and more^22–30^. Despite this essentiality, methionine restriction in an iso-caloric manner has been linked to increased insulin sensitivity and glucose metabolism^29^, more favorable cancer prognoses^31,32^, and lifespan extension^33–36^, however, the direct mechanisms of how low Met availability is sensed and signaled within the cell is not fully understood.

Met is directly converted into *S*-adenosylmethionine (SAM) via Methionine Adenosyltransferase I/II (MAT1/2A) in liver or exclusively via MAT2A in all remaining tissues^37^. Increasing the consumption of dietary Met is sufficient to increase SAM levels in human blood plasma, reflecting the dependency of SAM on cellular Met availability^38^. SAM abundance is impacted by both internal and external factors including circadian rhythm, age, metabolic input, disease state, cancer, and diet^19,39–41^. Interestingly, the availability of intracellular Met can drive reversible changes in methylated histones due to the intrinsic kinetic parameters of histone methyltransferases, including reversibly decreasing trimethylation upon restriction of Met^42,43^. Higher-order methylation states (i.e. me2/me3) have been proposed to act as reservoirs for methyl groups, thereby protecting nuclear SAM homeostasis and methylation potential^44–46^. This raises the possibility that specific epigenetic marks are preferentially preserved to maintain essential cellular functions, whereas others may serve as flexible reservoirs that store and release methyl groups in response to metabolic demand^19^.

Recent studies have shown that methionine depletion (Met-D) induces a conserved epigenetic adaptation characterized by a global loss of H3K9me2/me3 and active preservation of H3K9 monomethylation (H3K9me1). This response maintains heterochromatin integrity and supports epigenetic persistence following metabolic stress^42^. Mechanistically, H3K9 methyltransferases preferentially target unmethylated substrates under limiting SAM conditions, enabling *de novo* H3K9me1 deposition under nutrient deprivation^47^. However, most studies of methyl-metabolite restriction simultaneously deplete both Met and its downstream product SAM, making it unclear whether SAM depletion alone is sufficient to drive these epigenetic changes. Distinguishing the cellular responses to methionine versus SAM restriction is therefore critical for understanding how metabolic therapies may differentially impact epigenetic stability and for guiding strategies that selectively target tumor metabolism without compromising chromatin integrity.

Here, we directly tested whether intracellular SAM depletion is sufficient to drive the epigenetic adaptation observed during Met-D. Using pharmacologic inhibition and transient depletion of MAT2A to selectively reduce SAM without lowering intracellular Met, we compared the chromatin and transcriptional consequences of isolated SAM loss with those of Met-D. We further examined whether established SAM-responsive pathways contribute to this response by assessing the roles of SAMTOR and KDM4-family histone demethylases under nutrient stress. Our results show that selective SAM depletion fails to reproduce the heterochromatin remodeling, transposable element derepression, stress signaling, and transcriptional reprogramming induced by methionine deprivation. Instead, these findings identify Met availability itself as a primary metabolic signal that coordinates chromatin adaptation independently of methyl-donor abundance.

## Results

### MAT2B binds DNA in vitro but is dispensable for heterochromatin adaptation to methionine depletion

To investigate whether SAM synthesis is spatially regulated during Met-D, the behavior of the SAM synthetase, MAT2A, and its regulatory binding partner, MAT2B, was examined. MAT2B’s interaction (K_d_ = 6 ± 1 nM) is not required for MAT2A catalysis, but has been reported to increase MAT2A stability and catalytic abilities^37,48–50^. Previous reports noted that MAT2A/2B could co-localize with histone methyltransferases at gene promoters to quickly repress gene expression through direct SAM synthesis coupled with immediate deposition of H3K9me3 in iMEFs^51,52^. Therefore, we hypothesized that a similar coupled mechanism of locus-specific SAM synthesis could underlie the adaptive heterochromatin methylation observed under Met-D.

Following 24-hours Met-D, MAT2A protein levels increased in both cytoplasmic and nuclear fractions, with a ∼5-fold enrichment in the nucleus relative to ∼2-fold in whole-cell lysates (**Figure 1A**), indicating nuclear accumulation under nutrient stress. In contrast, MAT2B protein remained restricted to the cytoplasm and was undetectable in nuclear fractions under all conditions tested (**Figures 1B-C**). Consistent with this, MAT2B transcript levels were unchanged during Met-D, whereas MAT2A mRNA increased ∼30-fold (**Figure 1D**). These results demonstrate that catalytic MAT2A, but not regulatory MAT2B, is transcriptionally and spatially regulated under Met-D.

**Figure 1:**
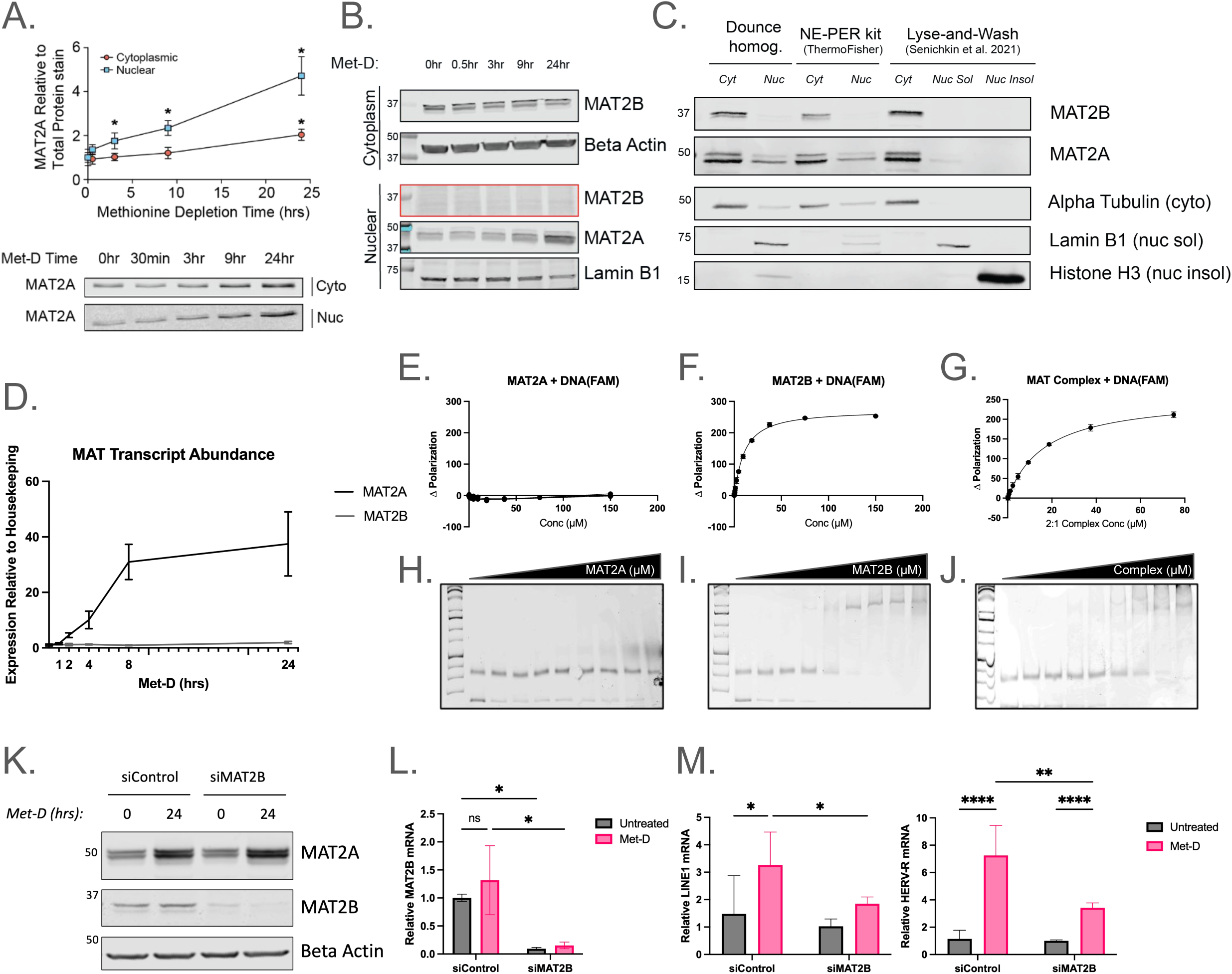
MAT2B binds chromatin *in vitro*, but is dispensable for heterochromatin adaptation to methionine depletion. (**A**) Above; Scatter plot of time-dependent increases in relative MAT2A abundance in HCT116 cells following Met-D. Below; Subcellular fractionation was performed on samples to separate cytoplasmic (cyto) versus nucleoplasm (nuc) fractions, which were then subjected to Western blot analysis. MAT2A abundance was determined relative to REVERT Total Protein Stain on the respective subcellular fraction. Error bars represent SD, n≥2, *p < 0.05 (Welch’s t-Test). (**B**) Western blot analysis of nuclear and cytoplasmic subcellular fractions under Met-D. MAT2B levels are stable within the cytosol but there is no indication of a translocation event, as seen with MAT2A nuclear accumulation. Lamin B1 was used as a marker for nucleoplasm, while Beta Actin was the cytosolic marker. (**C**) Three different subcellular fractionation techniques were used to assess nuclear MAT2B levels. Dounce homogenization is our standard in-house protocol, and was compared to NE-PER™ Nuclear and Cytoplasmic Fractionation Kit (ThermoFisher Scientific catalog #78833) and a published “Lyse-and-Wash” method^94^ which enables further separation of soluble and insoluble nuclear fractions. Insoluble nuclear fractions include mostly nuclear envelope proteins and proteins tightly bound to chromatin. Alpha Tubulin, Lamin B1, and Histone H3 were used as markers for cytoplasmic, soluble nuclear and insoluble nuclear fractions, respectively. Importantly, MAT2B was not observed in any nuclear fractionation technique. (**D**) Met-D time course RTqPCR analysis of MAT2A and MAT2B transcripts relative to internal housekeeping gene. n=3; error bars indicate SD. (**E-F**) Fluorescent Polarization (FP) analysis of varying concentrations of recombinantly purified enzyme (x axis) combined with 20bp of Widom 601 DNA labelled with fluorescein (FAM). Change in polarization indicates binding of the enzyme to DNA. (**G**) FP analysis of pre-incubated MAT2A and MAT2B in 2:1 ratio for complex formation prior to DNA-FAM addition. n=3; error bars represent SD. (**H-J**) Similar to FP experiments, increasing concentrations of recombinant MAT2A and MAT2B were incubated individually, and in 2:1 complex ratio, with recombinant nucleosomes before electrophoresis separation under non-denaturing conditions. Gels were stained with SYBR Safe DNA stain to visualize nucleosome-associated DNA. An upward migration of nucleosomes indicates a shift in molecular weight due to enzyme binding. n=1. (**K**) Western blot analysis of total MAT2A and MAT2B following a 48-hour siMAT2B knock-down followed by 24-hours of Met-D. MAT2B protein levels decrease by ∼ 80%, relative to Beta Actin loading control. MAT2A levels are unaffected by the loss of MAT2B. Representative image. (**L**) RTqPCR analysis of MAT2B transcripts relative to internal housekeeping gene, following the experimental design in (K). n=3; error bars represent SD; *p value < 0.05 (2way ANOVA). (**M**) RTqPCR analysis of TE expression levels following 48-hours siMAT2B and 24-hours of Met-D. n=3; error bars represent SD; *p value < 0.05 (2way ANOVA).

We next tested whether MAT2A or MAT2B can directly associate with chromatin components. Fluorescence polarization assays showed that recombinant MAT2A did not bind free DNA (**Figure 1E**). In contrast, MAT2B bound DNA with low affinity (K_d_ ∼13.5 µM), and pre-formed MAT2A–MAT2B complexes exhibited similar micromolar binding (**Figures 1F-G**). Importantly, the order of magnitude difference between MAT2B’s reported K_d_ for its binding partner (MAT2A) compared to its observed K_d_ for DNA suggest that MAT complex formation precedes and persists while MAT2B interacts with DNA. Electrophoretic mobility shift assays further demonstrated that MAT2B, but not MAT2A, binds recombinant nucleosomes *in vitro* (K_d_ ∼5.1 µM), and this interaction was retained in the presence of MAT2A (**Figures 1H-J**).

Despite this *in vitro* binding capacity, MAT2B did not localize to the nucleus under Met-D (**Figures 1B-C**), suggesting chromatin-associated SAM synthesis in the cell conditions described here are unlikely. To directly test whether MAT2B contributes to heterochromatin regulation, we depleted MAT2B by siRNA prior to Met-D. MAT2B knockdown (>90% transcript and >80% protein depletion) importantly did not alter MAT2A protein levels (**Figures 1K-L**). Next, RTqPCR was used to probe the transcript levels of transposable elements (TEs) as a proxy for heterochromatin adaptation (i.e. de-repression) under Met-D^42^. Notably, MAT2B depletion reduced transposable element (TE) expression under Met-D (**Figure 1M**), indicating that MAT2B is not required for heterochromatin maintenance and may modestly oppose TE repression under these conditions.

Together, these data show that although MAT2B can bind chromatin substrates *in vitro*, this interaction is not sufficient for its recruitment *in vivo* and MAT2B is not required for heterochromatin adaptation to methionine depletion. Instead, these findings indicate that nuclear accumulation of MAT2A, rather than MAT2B-mediated chromatin targeting, likely contributes to SAM-dependent chromatin regulation during Met-D.

### MAT2A inhibition depletes intracellular SAM without reducing methionine levels

Nuclear accumulation of MAT2A under Met-D suggests a mechanistic link between nuclear SAM synthesis and epigenetic adaptation, therefore, the role of active SAM synthesis was examined next. AG-270 is a recently developed small-molecule allosteric inhibitor of MAT2A (IC_50_ = 14 nM) with promising results in Phase I clinical trials for advanced malignancies lacking MTAP, a key enzyme in the methionine salvage pathway^53,54^. To determine whether SAM depletion alone is sufficient to mimic Met-D, we pharmacologically inhibited MAT2A using AG-270 and compared the metabolic effects with those of Met-D.

First, to test the metabolic effects of MAT2A pharmacological inhibition on whole-cell methionine and SAM levels, a dose-response analysis of varying concentrations of AG-270 was performed alongside Met-D in HCT116 cells (**Figure 2A**). Metabolites were extracted after 24-hours and processed by liquid chromatography-tandem mass spectrometry (LC-MS/MS) to determine relative metabolite abundance normalized to total protein. LC-MS/MS analysis revealed that Met-D drastically reduced both intracellular methionine and SAM, as expected (**Figure 2B**). In contrast, AG-270 treatment selectively depleted SAM in a dose-dependent manner without decreasing methionine levels. Notably, 1 µM AG-270 treatment reduced SAM abundance to levels comparable to Met-D while preserving intracellular methionine.

**Figure 2:**
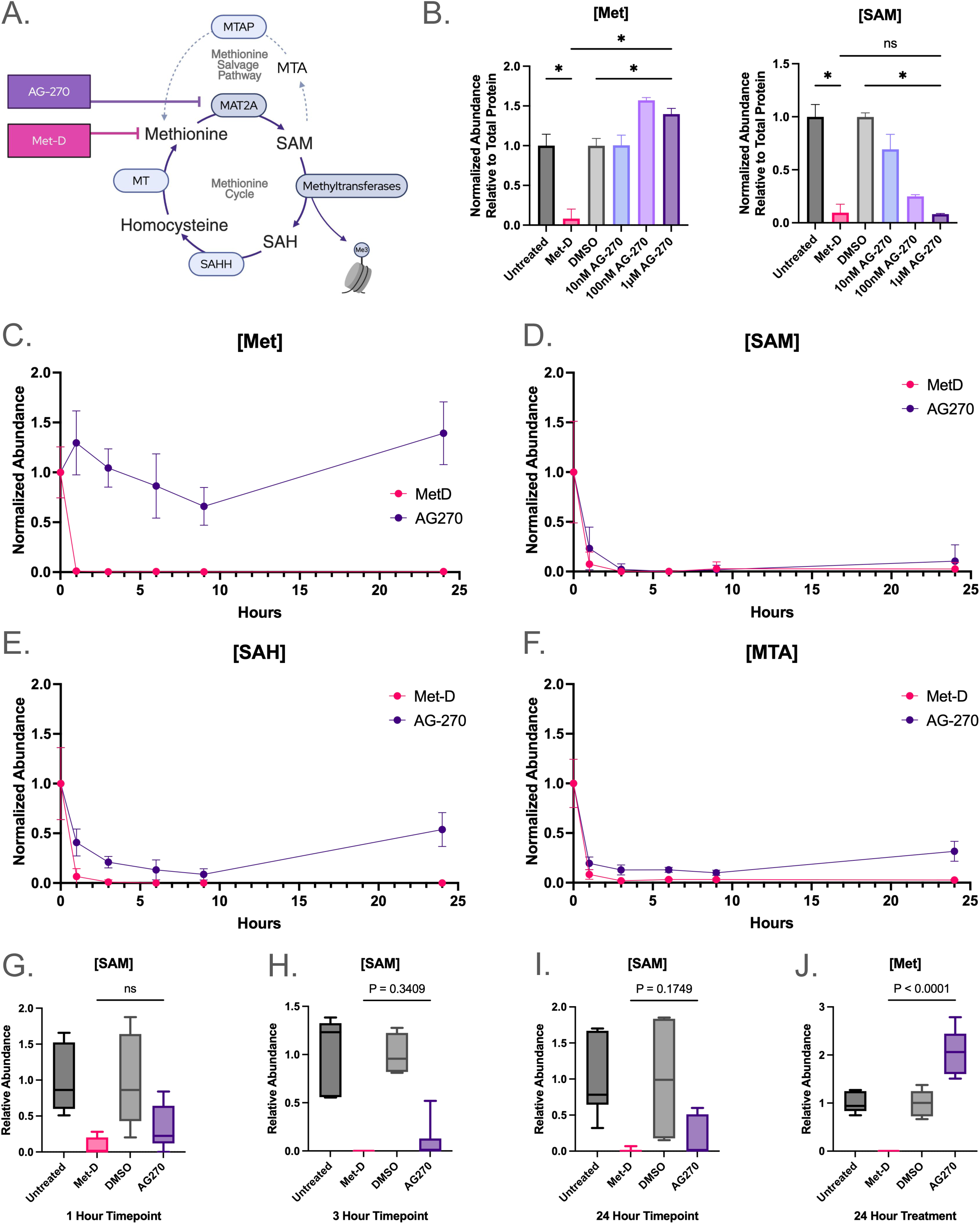
MAT2A inhibition with AG-270 depletes intracellular SAM levels without negatively affecting methionine content. (**A**) Metabolic diagram of the methionine cycle with experimental perturbations; dietary methionine depletion (Met-D, pink) and MAT2A inhibition with AG-270 (purple). Enzymes are shown in ovals, metabolites in black text. (**B**) Metabolite abundance relative to appropriate control following 24-hours of growth in methionine-depleted medium or complete medium containing varying concentrations of AG-270, dissolved in DMSO. Importantly, 1µM AG-270 treatment depletes SAM to comparable levels as Met-D, without limiting Met. n=3; error bar represents SD; *p<0.05 (Ordinary one-way ANOVA). (**C-F**) Metabolite abundance, normalized to internal untreated time=0 control, following Met-D and 1µM AG-270 treatment with metabolites extracted at 1, 3, 6, 9, and 24-hours. n=6; error bar represents SD. (**G-I**) SAM or (**J**) Met abundance, relative to time-matched treatment control, at 1, 3, 24-hour time points. n=6; p values comparing Met-D and AG-270 as indicated (Welch’s t-Test).

Next, the kinetics of metabolite depletion were compared. Under Met-D, intracellular Met is rapidly depleted within 5 minutes, while SAM concentration consequently reaches undetectable levels within 1-hour^55^. The timing of metabolite depletion under Met-D is critical to allow visualization of downstream epigenetic changes within 24-hours. Methionine levels were rapidly depleted under Met-D but remained stable during AG-270 treatment, as expected (**Figures 2C,J**). In contrast, SAM levels declined rapidly under both conditions, reaching near-complete depletion within hours and remaining suppressed through 24-hours (**Figures 2D,G–I**). Importantly, SAM levels were indistinguishable between Met-D and AG-270 treatment at early time points and fell below detection limits by 3-hours in both conditions (**Figures 2G–H**). Consistent with reduced SAM availability, divergent downstream metabolites *S*-adenosylhomocysteine (SAH) and methylthioadenosine (MTA) were decreased under both treatments (**Figures 2E-F**).

These results demonstrate that pharmacological inhibition of MAT2A recapitulates the magnitude and kinetics of SAM depletion observed during Met-D, while maintaining intracellular methionine levels. These conditions, therefore, establish a system in which SAM depletion can be uncoupled from methionine availability. Next, we used this system to determine whether isolated SAM depletion is sufficient to drive the chromatin and transcriptional changes observed under Met-D.

### SAM depletion alone does not induce heterochromatin adaptation

To determine whether loss of SAM is sufficient to drive heterochromatin remodeling, the effects of Met-D and pharmacological SAM depletion (AG-270) were compared. Met-D elicits a conserved loss of H3K9me3/me2 and a concurrent upregulation of TEs such as LINES and HERVs, likely due to the loss of repressive PTMs^16,42^. To investigate the effects of metabolite depletion on heterochromatin, transposable element (TE) derepression was used as a proxy for heterochromatin instability via lost epigenetic repression.

Met-D robustly induced TE expression, including LINE1 and multiple HERV families (**Figures 3A–B**). In contrast, SAM depletion to equivalent levels using AG-270 did not increase TE expression, indicating that equivalent depletion of SAM is not sufficient to destabilize heterochromatin. This phenotype was reproduced in HEK293T cells, demonstrating consistency across cell types (**Figure S1**).

**Figure 3:**
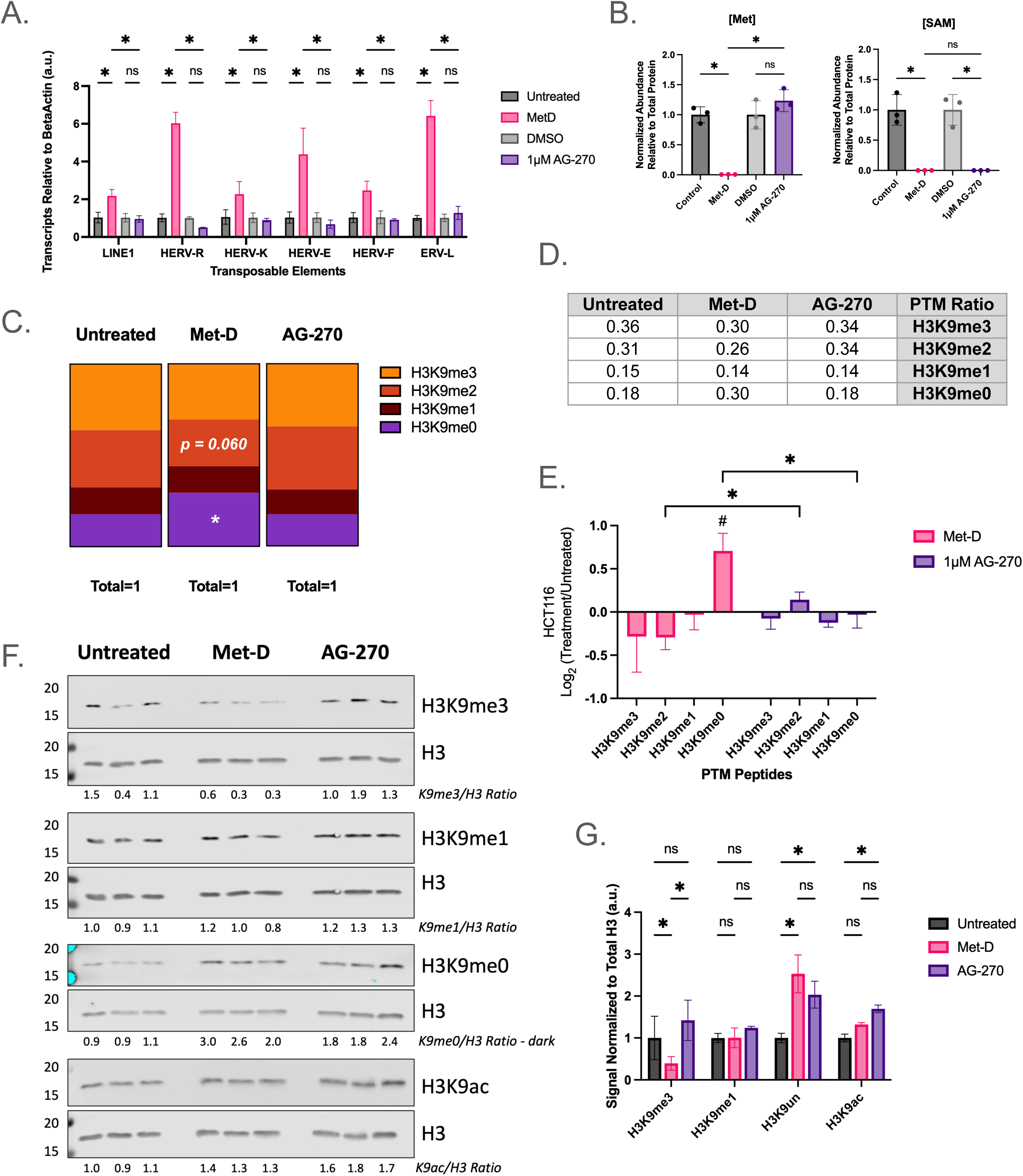
SAM depletion alone does not induce heterochromatin adaptation. (**A**) Bar graph depicting transcript abundance of heterochromatic transposable elements relative to internal housekeeping gene and normalized to the appropriate DMSO-matched control following 24-hours of indicated treatment. n=3 biological replicates; error bars represent SD; *p < 0.05 (2way ANOVA). (**B**) LC-MS/MS metabolite abundance of intracellular Met and SAM relative to appropriate control following 24-hours of growth in Met-D or 1µM AG-270 treated medium. n=3; error bars represent SD; *p < 0.05 (Ordinary one-way ANOVA). (**C**) Stacked bar graphs depicting normalized ratio of H3K9-methyl PTM relative to total H3K9 peptide (residues 9-17) abundance derived via LC-MS/MS. The sum of all segments within a condition is 1. n≥2; *p < 0.05 (Welch’s t-Test). (**D**) Chart depicting relative ratios of PTMs from (C). (**E**) Bar graph illustrating LC-MS/MS-generated log2 fold-changes for individual H3K9 PTMs. n≥2. (**F**) Western blot analysis of PTM abundance relative to internal total H3 signal, performed on acid-extracted histones following 24-hours of Met-D or 1µM AG-270 treatment. PTM / Total H3 was determined within each biological replicate (n=3) and normalized to averaged ratio for untreated conditions. (**G**) Plotted normalized ratios from (F). n=3; error bars represent SD; *p < 0.05 (Welch’s t-Test).

Next, histone H3K9 methylation states were examined by LC-MS/MS. Met-D reduced nuclear H3K9me2/me3 while maintaining H3K9me1, consistent with prior observations^42^ (**Figures 3C–D**). In contrast, AG-270 treatment largely preserved the distribution of H3K9 methylation states, with only minor, non-significant changes. Analysis of log2 fold changes confirmed that Met-D induces substantially greater shifts in H3K9 methylation than SAM depletion alone (**Figure 3E**). These findings were validated by quantitative Western blot analysis (**Figures 3F-G**).

Together, these results demonstrate that Met-D, but not depletion of SAM alone, induces heterochromatin remodeling characterized by loss of higher-order H3K9 methylation states and transposable element derepression. These data establish that depletion of the methyl-donor SAM is not sufficient to trigger the heterochromatin adaptation response to Met-D, therefore, we next sought to identify the mechanisms that distinguish methionine depletion from isolated SAM depletion.

### SAMTOR is dispensable for heterochromatin adaptation to methionine depletion

To date, no direct cellular sensors of free methionine have been identified. In contrast, SAMTOR has been reported as a direct SAM-sensing protein responsible for impeding growth signals under low SAM availability through inhibition of the master cell regulator, mechanistic target of rapamycin complex 1 (mTORC1)^56–58^. SAMTOR, through its negative regulation of mTORC1, is purported to act as a cellular “brake” to turn off proliferation and metabolism under SAM starvation until the stress can be resolved (**Figure S2A**). Therefore, if SAMTOR’s SAM-sensing properties are required for epigenetic regulation under Met-D, then loss of SAMTOR itself would lead to a further derepression of TEs, potentially through maintained mTORC1 signaling.

To test whether SAM sensing via SAMTOR mediates epigenetic adaptation to Met-D, SAMTOR was acutely knocked down using siRNAs prior to Met-D. Efficient depletion (∼91% mRNA reduction) was confirmed by RT-qPCR (**Figure S2B**). However, SAMTOR knockdown did not increase LINE1 or HERV-R expression under Met-D (**Figures S2C-D**), indicating that SAMTOR is not required for TE derepression. Consistent with this, phosphorylation of S6K revealed that mTORC1 signaling remained inactive after 24-hours of Met-D, regardless of SAMTOR status (**Figure S2E**). This observation agrees with prior reports that mTORC1 activation is largely resolved by later stages (i.e. 24-hours) of amino acid deprivation^59,60^.

Together, these results demonstrate that SAMTOR and mTORC1 signaling are dispensable for heterochromatin remodeling at late stages of Met-D, indicating that SAM sensing through this axis does not mediate the epigenetic response.

### KDM4 demethylases contribute to, but are not required for, Met-D–induced heterochromatin adaptation

Depletion of the methyl donor SAM alone failed to recapitulate the heterochromatin adaptation observed under Met-D, which was unexpected given the central role of SAM in histone methylation. Our findings further argue that canonical SAM-sensing pathways are dispensable for this heterochromatic response. This disconnect suggests that Met-D–induced epigenetic remodeling is not driven solely by reduced methyl-donor availability, but instead requires additional methionine-dependent signals.

Under Met-D, maintenance of H3K9me1 depends on *de novo* monomethylation reactions^42^, likely supported by the intrinsic catalytic activity of H3K9 methyltransferases^47^. For this adaptive redistribution of methylation states to occur, higher-order methyl marks (H3K9me2/3) must first be removed to generate unmethylated substrates (H3K9un or H3K9me0). The KDM4 family (KDM4A–D) of histone demethylases, which catalyze removal of H3K9me2/3^61^, therefore represents a candidate mediator of this process and a potential point of divergence between Met-D and SAM depletion.

RT-qPCR analysis revealed that Met-D, but not SAM depletion via AG-270, significantly increased KDM4A–D transcript levels in HCT116 and HEK293T cells (**Figure 4A**, **Figure S1C**), suggesting a methionine-specific induction of demethylase expression. Previous reports link serine-depletion dependent KDM4C activation to the demethylation of gene promoters and subsequent transcription of Activating Transcription Factor 4 (ATF4), a key enzyme in the Integrated Stress Response (ISR)^62^. Consistent with a broader stress response, ATF4 expression was strongly induced by Met-D but not AG-270 (**Figure 4B**). MAT2A transcript and protein levels increase significantly under both treatments (**Figure 4C**), consistent with the SAM-dependent regulation of MAT2A mRNA stability^40,63,64^.

**Figure 4:**
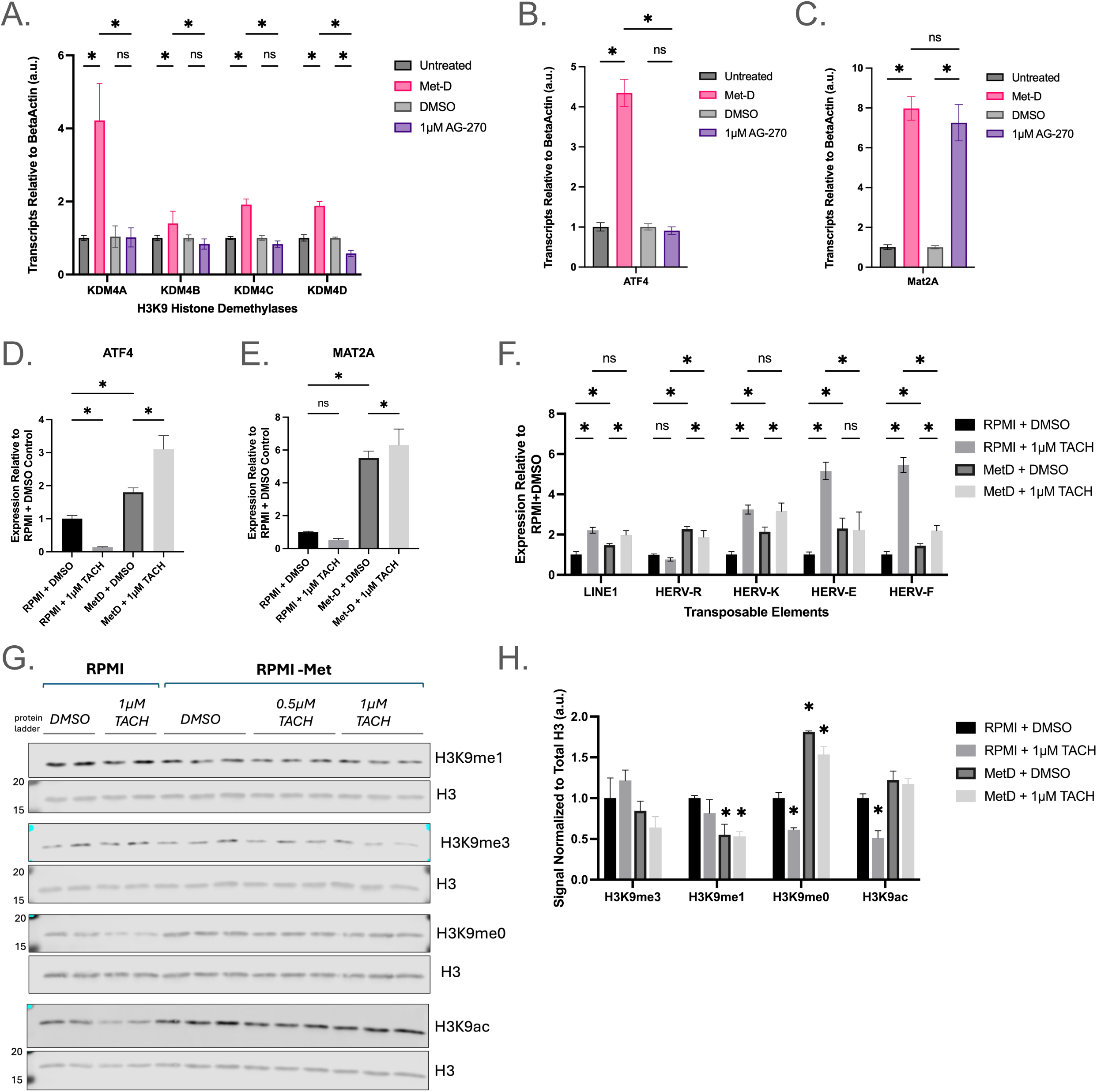
KDM4 demethylases contribute to, but are not required for, Met-D-induced heterochromatin adaptation. (**A**) Bar graph depicting transcript abundance of KDM4 family of H3K9me2/me3 demethylases relative to internal housekeeping gene and normalized to the appropriate DMSO-matched control following 24 hours of indicated treatment. n=3 biological replicates; error bars represent SD; *p < 0.05 (2way ANOVA). (**B-C**) Relative mRNA abundance of stress-induced ATF4 and the SAM-synthetase MAT2A, as in (A). (**D-E**) Relative mRNA abundance of MAT2A (D), ATF4 € and transposable elements (F) following KDM4 inhibition with 1µM TACH101 combined with Met-D. HCT116 adherent cells were pre-treated for one hour with complete medium (RPMI) containing TACH101 or DMSO. After one hour, cells were given fresh methionine-complete or -depleted media, containing either DMSO control or TACH101. Cells were harvested following 24 hours of incubation. (**G**) Western blot analysis of PTM abundance relative to internal total H3 signal, performed on acid-extracted histones following 24 hours of indicated medium combined with TACH101. (**H**) Bar graph depicting quantified PTM / Total H3 abundance from (J), normalized to RPMI + DMSO control. n=2-3; error bars represent SD. *p value < 0.05 (Welch’s t-Test).

To test whether KDM4 activity is required for Met-D-induced adaptation, cells were treated with the pan-KDM4 inhibitor, TACH101, prior to and throughout Met-D. TACH101, or Zavondemstat, is a small molecule pan-inhibitor that specifically targets KDM4 family members A-D (IC_50_ ≤ 0.080 µM *in vitro*) with minimal inhibition of other KDM families^65,66^.

Contrary to expectation, KDM4 inhibition did not suppress Met-D-dependent stress signals. Instead, both MAT2A and ATF4 were further increased under combined treatments (**Figures 4D-E**). While KDM4C activity may contribute towards ATF4 induction in a Met-deprivation context similar to reports under serine limitation^62^, KDM4 activity is not required for ATF4 expression. These results contradict the original hypothesis, which predicted that blocking demethylation would dampen cellular stress responses.

Consistent with these results, combined KDM4 inhibition and Met-D treatment further increased the expression of multiple transposable elements (LINE1, HERV-K, HERV-F), while others (HERV-R, HERV-E) remained elevated without additional induction (**Figure 4F**). Histone analysis by quantitative Western blots confirmed effective KDM4 inhibition, as evidenced by increased H3K9me3 (∼20%) under control conditions; however, Met-D–induced loss of H3K9me3 persisted and was further exacerbated by TACH101 (**Figures 4G–H**). These data is counter to the expected role of KDM4 enzymes and argues that H3K9me3 loss is not primarily driven by enzymatic demethylation, suggesting alternative mechanisms such as compensatory demethylase activity, impaired methyltransferase dynamics or increased histone turnover^43,67^.

Together, these findings demonstrate that although KDM4 expression is induced during Met-D, KDM4 catalytic activity is not required for reduced levels of H3K9me3, transposable element derepression, or ATF4 activation. Instead, KDM4s appear to modulate, but not drive, the epigenetic response to Met-D. Consistent with this, inhibition of SAM synthesis via AG-270 fails to induce KDM4 expression or downstream epigenetic remodeling, further supporting that loss of SAM alone is insufficient to trigger this adaptive program.

### Methionine depletion activates an innate immune response consistent with viral mimicry

Given the strong induction of TE expression under Met-D, we asked whether this triggers an innate immune response via endogenous double-stranded RNA. It is well documented that perturbation of heterochromatin or heterochromatin-modifying enzymes can exacerbate TE expression, thereby forming harmful double-stranded nucleic acids which mimic viral infection and activate integrated stress pathways ^68–71^.

RT-qPCR analysis of interferon-stimulated genes (ISGs) and nucleic acid sensing pathways revealed robust activation under Met-D, but not SAM depletion (**Figure S3**). Notably, IFNA1 (∼30-fold increase), and cGAS, and IRF7 (∼11-fold increase each) were strongly upregulated, along with additional ISGs. These results are consistent with, but not sufficient to establish, a model in which TE-derived nucleic acids activate innate immune pathways. Future studies are needed to determine whether TE-derived dsRNA directly activates PKR or cGAS-STING signaling during methionine stress. Thus, Met-D uniquely induces an innate immune transcriptional program that is not recapitulated by SAM depletion alone.

### Methionine depletion elicits transcriptional programs distinct from SAM limitation

To define global transcriptional responses, paired-end non-directional Illumina-based mRNA sequencing following Met-D and AG-270 treatments was performed. As a note, TE transcripts are not quantified under traditional RNA-seq pipelines, largely due to their repetitive nature which requires long-read sequencing^72^.

Replicates showed high reproducibility (R² ≥ 0.98), and 23,000 genes were detected across conditions (**Figure 5A**). Principal component analysis (PCA) of normalized gene expression values (i.e. FPKM) exhibited distinct profiles for both treatments compared to controls, while z-score clustering analysis revealed a highly distinct transcription profile under Met-D (**Figures 5B-C**). Differential expression (DE) analysis identified ∼4,500 genes altered by Met-D, compared to only 68 genes with AG-270 (**Figures 5D-F**), with 37 differentially expressed genes (DEGs) shared (**Figure 5G, Table S5**). Reactome analysis^73^ of the shared DEGs revealed slight enrichment for YAP1-mediated gene expression (**Table S6).** Consistent with earlier results, ATF4 was strongly induced by Met-D but not AG-270, despite increased MAT2A expression in both conditions (**Figures 5H-I**).

**Figure 5:**
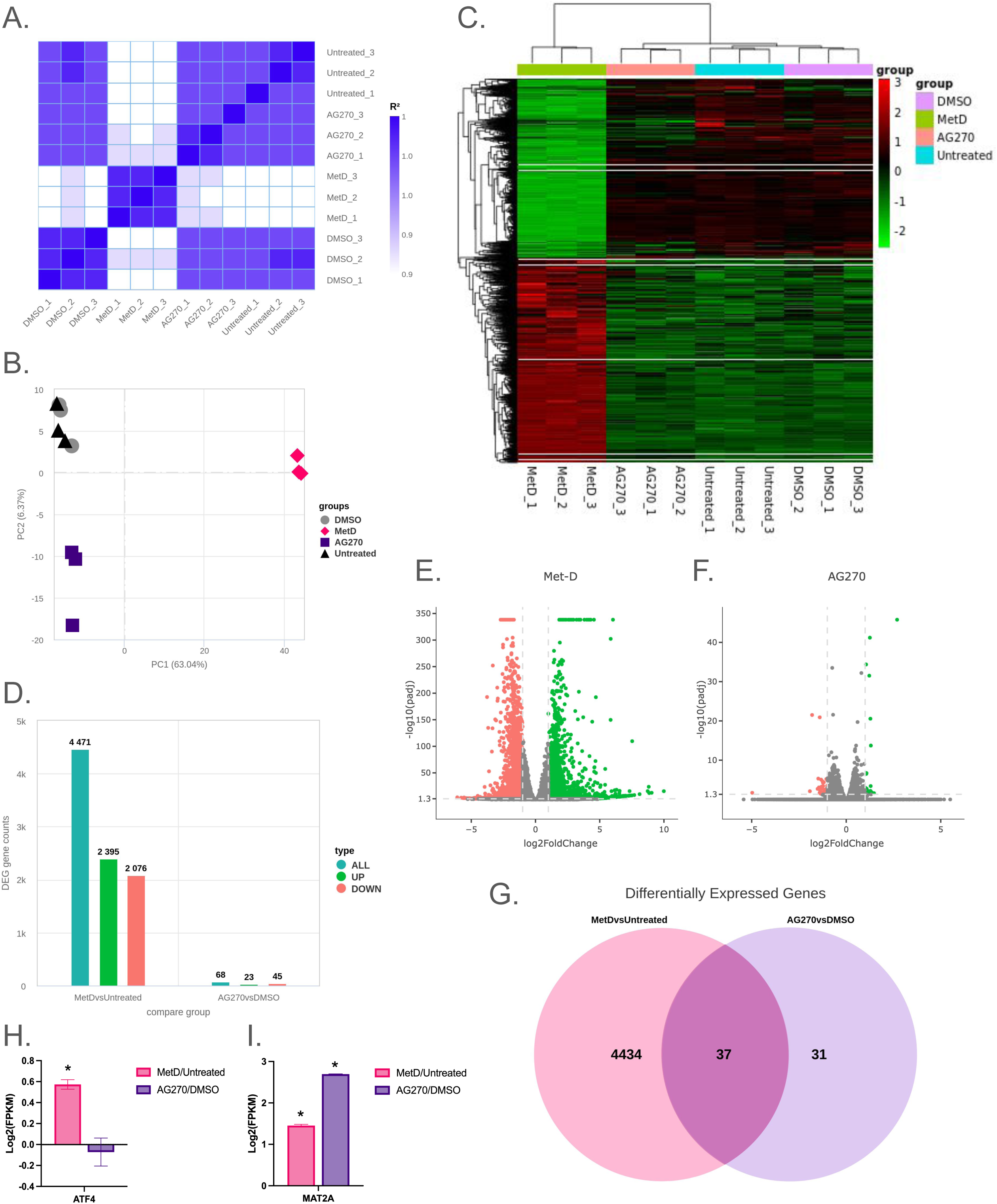
Methionine depletion elicits transcriptional programs distinct from SAM limitation. (**A**) Heat map of pairwise Pearson correlation coefficients (R^2^) computed from normalized gene expression values across Met-D and AG-270, as well as appropriate controls. The closer the correlation coefficient is to 1 the higher similarity between samples (**B**) Plotted principal component analysis (PCA) on gene expression values (FPKM). (**C**) Heatmap of Z-score-normalized differentially expressed genes compared to a pooled differential gene set. Colors reflect relative gene expression compared to all samples. Hierarchical clustering was applied to the FMPK values of genes homogenized by Z-score, where genes with similar expression patterns will be clustered together on the left, while clustering on the top of the graph represents similarities in gene expression between samples. (**D**) Bar graph depicting number of differentially expressed genes identified for each treatment relative to indicated control, separated by the number of total, up-regulated, and down-regulated differentially expressed genes. (**E-F**) Volcano plots of significant differentially expressed genes for indicated treatments. The x-axis represents fold-change in gene expression between samples, and y-axis indicates the statistical significance of the differences. Each dot represents a gene, while red indicates significant downregulation and green indicates significant upregulation. (**G**) Venn diagram depicting shared and unique differentially expressed genes among treatments. (**H-I**) Bar graphs depicting relative expression (FPKM) of ATF4 and MAT2A in each treatment compared to indicated control. n=3; error bars indicate SD; *p value <0.05 (Welch’s t-Test).

Gene Set Enrichment Analysis (GSEA) for DE transcripts in treatments relative to controls was performed. For Met-D treated cells, N-Acetyltransferase Activity (GO:0008080) was the only Gene Ontology (GO) pathway was significantly enriched with an FDR cut-off q < 0.05 (**Figure 6A**). Increasing the FDR to q > 0.15 identified over 2,000 GO enrichments including positive regulation of defense mechanisms and stress-related transcription, while phosphate metabolism was negatively regulated (**Figure 6B**). KEGG (Kyoto Encyclopedia of Genes and Genomes) and Reactome analysis further supported these data, with high enrichment for pathways involved in metabolic rewiring, cell cycle and replicative arrest, structural alterations, and various signaling pathways (**Figures 6C-E**). In stark contrast, AG-270 revealed no significantly altered pathway enrichments in GSEA, KEGG, or GO analyses (**Figures 6F-I**). Affected biological processes were widespread and included ErbB, EGFR, and Wnt signaling pathways, as well as structural organization. These data support the hypothesis that Met-D stimulates a drastic transcriptional program spanning multiple signaling pathways and metabolic processes, while equivalent reduction of SAM alone exhibited minimal effects on transcriptional programs.

**Figure 6:**
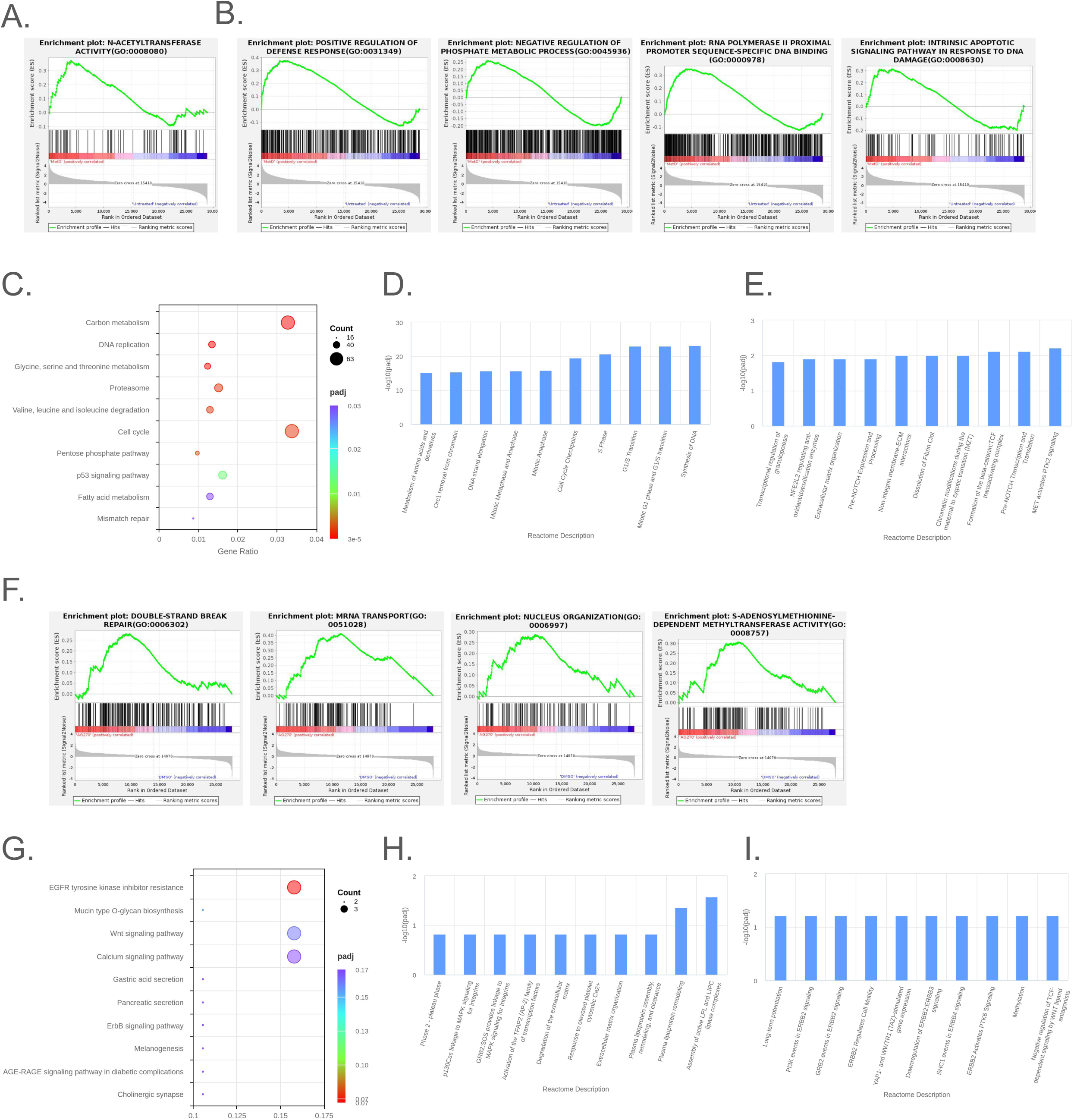
Pathway enrichment analysis of nutrient manipulations in HCT116 cells. (**A-B**) Gene set enrichment analysis (GSEA) of Met-D specific GO terms plotted as enrichment scores (green line) with an FDR cut-off of q<0.05 (A) or q<0.15 (B). (**C**) KEGG enrichment analysis of pathways altered by Met-D. Bubble colors indicate p-adj and size indicates the number of differentially expressed genes identified within the KEGG pathway. (**D**) Reactome pathway analysis of down-regulated processes under Met-D. For pathway enrichment, Reactome uses a threshold of padj < 0.05 to determine significant enrichment. (**E**) Reactome enrichment of up-regulated processes under Met-D as in (D). (**F**) GSEA of non-significant GO terms under AG-270 treatment as in (B). (**G**) KEGG enrichment plot for altered pathways in AG-270-treated cells. (**H-I**) Reactome enrichment pathways for down-regulated (H) and up-regulated (I) processes under AG-270.

Collectively, these data demonstrate that methionine depletion specifically triggers a robust and distinct transcriptional program that is not recapitulated by depletion of SAM.

### MAT2A knockdown reduces SAM but produces a mixed heterochromatin phenotype and distinct transcriptional outcomes

To test the effects of SAM depletion through a non-pharmaceutical approach, MAT2A was transiently knocked down in HCT116 cells, effectively reducing SAM levels by ≥80% without altering methionine abundance (**Figure S4A**). Reduction in MAT2A RNA (≥91%) and protein levels (≥72%) were observed by RTqPCR and Western blot analysis, respectively, after 48- and 72-hours knock-down (**Figures S4B-D**). Despite substantial SAM reduction, TE expression was not uniformly increased. LINE1 and HERV-R were significantly repressed, whereas HERV-K and HERV-F were modestly induced (**Figure S4E**), indicating a mixed heterochromatin response. Expression of various cellular Met-D stress sensors (i.e. ATF4 and innate immunity sensors) and KDM4 family members were concordantly mixed, though majority were uninduced (**Figure S4F-H**).

The siMAT2A transcriptome was assessed and appeared to reflect a secondary stress-adaptation program rather than a discrete “low-SAM” signature, with 5,388 total differentially expressed genes identified and nearly equal numbers of upregulated and downregulated gene expression changes (**Table S5, Figure S4I**). The most internally consistent pattern was the activation of the integrated stress response (ISR), supported by the highly significant (*p*adj << 0.001) induction of canonical ATF4 target genes DDIT3/CHOP (Log2FC :: 1.0) and TRIB3 (Log2FC :: 1.1), alongside increased CDKN1A/p21 (Log2FC :: 2.1), which often accompanies ISR-associated growth arrest^74–76^. In parallel, strong induction of EGR1, EGR3, and KLF15 suggests engagement of a broader immediate-early transcriptional response, implying that loss of MAT2A influences signaling networks beyond methyl-donor metabolism^77^. Following KEGG and Reactome analysis of siMAT2A-specific DEGs, steroid biosynthesis and SREBP-associated cholesterol metabolism was shown to be altered (**Figures S4K-M**), consistent with many SREBP target genes (i.e. FASN, SCD, ACACA, HMGCR, LDLR, and INSIG1) showed altered expression profiles (**Table S5**). While a direct connection between MAT2A levels and SREBP function has yet to be established, SREBP-dependent lipid biosynthetic programs are known to respond to nutrient stress and chromatin-dependent transcriptional regulation^78,79^.

This transcriptional rewiring suggests that prolonged MAT2A loss perturbs chromatin accessibility or transcriptional fidelity independently of bulk SAM reduction. Furthermore, these data suggest that while MAT2A knockdown reduces SAM, it triggers a mixed stress-state with ectopic transcriptional remodeling rather than the simple methylation-dependent response observed under Met-D. Notably, siRNA-mediated knockdown can independently activate innate immune and stress pathways through cellular sensing of double-stranded RNA and transfection-associated effects, potentially contributing to the mixed stress phenotype observed in siMAT2A cells^80,81^. Given the pleiotropic roles of MAT2A in cellular physiology, including effects on translation and transcription independent of methylation, these results suggest that direct SAM reduction via MAT2A knock-down does not phenocopy Met-D, nor AG-270-stimulated SAM depletion. Unlike acute pharmacologic inhibition, siRNA-mediated depletion likely induces chronic metabolic imbalance and compensatory transcriptional rewiring, obscuring direct effects of SAM depletion.

## Discussion

Methionine availability is tightly linked to epigenetic regulation through conversion into S-adenosylmethionine (SAM), the universal methyl donor for histone and DNA methylation. Beyond serving as a substrate for protein synthesis, methionine occupies a central position in one-carbon metabolism, where it contributes to redox balance, nucleotide biosynthesis, and methylation-dependent gene regulation^22,23,27^. Through conversion to SAM, methionine directly fuels histone and DNA methyltransferases, thereby coupling metabolic state to chromatin structure and gene expression^19,42,43,45^. Through this role, methionine functions as a metabolic node that dynamically couples nutrient availability to chromatin state. Although changes in methionine availability have long been known to alter cellular physiology, the extent to which these responses are driven by methionine versus its downstream methyl-donor SAM has remained unresolved.

This question is particularly important considering the broad physiological consequences of methionine restriction. Dietary methionine limitation improves metabolic health, reduces inflammation, and extends lifespan in multiple model systems^29,34,36,82–86^. In parallel, cancer cells exhibit a well-documented dependence on exogenous methionine—termed the Hoffman effect—where methionine limitation selectively impairs tumor proliferation while sparing normal cells^24^. Consistent with this vulnerability, elevated expression of the SAM-synthesizing enzyme MAT2A and its regulatory subunit MAT2B, as well as aberrant nuclear localization of these proteins, have been associated with aggressive disease and poor clinical prognosis across multiple cancer types, underscoring the importance of SAM synthesis and compartmentalization in tumor biology^87,88^. These observations suggest that dysregulation of methionine metabolism is not merely a metabolic adaptation but is tightly linked to epigenetic control mechanisms that support malignancy. Despite these robust phenotypic outcomes, the molecular mechanisms underlying methionine sensing and adaptation, particularly at the level of chromatin regulation, remain poorly defined. Moreover, while dietary methionine restriction has shown promise, its clinical implementation is limited by poor long-term compliance, highlighting the need for alternative strategies that recapitulate the beneficial effects^89,90^.

Here, we show that methionine depletion (Met-D) elicits a robust and coordinated epigenetic and transcriptional response that is not reproduced by selective depletion of SAM alone. Although pharmacologic inhibition of MAT2A or transient MAT2A knockdown reduced intracellular SAM to levels comparable to Met-D, neither approach induced the characteristic redistribution of H3K9 methylation, transposable element (TE) derepression, or widespread transcriptional responses observed under methionine restriction. Instead, isolated SAM depletion was largely tolerated despite near-complete loss of the methyl donor. These findings challenge the prevailing view that SAM availability is the primary determinant of methylation-dependent chromatin regulation and instead identify methionine as a critical upstream signal governing epigenetic adaptation.

A central observation of this study is the dissociation between intracellular SAM abundance and heterochromatin remodeling. Met-D caused the expected loss of H3K9me2 and H3K9me3 together with preservation of H3K9me1 at these loci, consistent with prior evidence that cells preferentially maintain lower-order methylation states when methyl donors become limiting^42,47^. At the same time, Met-D induced TE expression, indicating that this redistribution only partially preserves heterochromatin function. The selective retention of H3K9me1 at these loci may serve as a molecular memory of previously repressed chromatin, thereby preserving a permissive substrate that can be rapidly restored to higher methylation states when methionine availability is re-established^42,47^. Thus, the adaptive methylation response appears sufficient to maintain some aspects of chromatin organization while failing to fully suppress repetitive element transcription during prolonged nutrient stress. In contrast, SAM depletion alone did not significantly alter H3K9 methylation or TE repression, indicating that reduced methylation capacity alone is insufficient to trigger this broader epigenetic program. This dissociation between methyl-donor abundance and chromatin remodeling suggests that enzymatic methylation capacity is not the rate-limiting determinant of epigenetic adaptation under nutrient stress.

The observed nuclear accumulation of MAT2A during Met-D further supports the idea that cells actively respond to methionine stress. MAT2A protein increased in both whole-cell and nuclear fractions, whereas MAT2B remained cytoplasmic despite its ability to bind DNA and nucleosomes *in vitro*. These findings argue against a model in which MAT2B recruits MAT2A directly to chromatin under these conditions. Instead, nuclear enrichment of MAT2A may represent a compensatory mechanism to preserve local SAM synthesis in the nucleus during nutrient limitation. However, because direct depletion of SAM failed to reproduce chromatin phenotypes, nuclear MAT2A accumulation alone is unlikely to be sufficient to initiate the adaptive response.

Our results further clarify that the cellular response to methionine deprivation is fundamentally distinct from the consequences of directly perturbing SAM synthesis. While pharmacologic inhibition of MAT2A with AG-270 produced minimal transcriptional changes, consistent with the limited epigenetic remodeling observed under SAM depletion alone, siRNA-mediated depletion of MAT2A triggered a secondary stress-adaptation program. This response was characterized by activation of the Integrated Stress Response (ISR), including induction of canonical stress markers such as DDIT3 and TRIB3, indicative of proteotoxic or nutrient stress. Importantly, siMAT2A fails to recapitulate the robust innate immune activation and coordinated transposable element derepression observed under Met-D, reinforcing the conclusion that methionine availability itself—rather than the absolute abundance of SAM—acts as the primary metabolic signal coordinating integrated epigenetic and transcriptional adaptation to nutrient stress. This partial stress state suggests that MAT2A depletion elicits indirect consequences beyond loss of catalytic activity, potentially reflecting non-catalytic roles for MAT2A or chronic metabolic imbalance that engages compensatory pathways, including feedback attenuation of receptor tyrosine kinase signaling (via *ERRFI1*) and oxidative stress responses (via *FTL*). The enrichment of SREBP-associated lipid metabolic genes after MAT2A depletion raises the possibility that chronic suppression of SAM synthesis perturbs membrane or sterol homeostasis, although whether this reflects a direct consequence of altered methylation capacity or a secondary stress adaptation remains unclear.

Mechanistically, our data argue against a primary role for canonical SAM-sensing pathways in mediating epigenetic remodeling under Met-D. SAMTOR has been proposed to sense intracellular SAM and regulate mTORC1 signaling under methyl-donor limitation^56^. However, depletion of SAMTOR did not alter TE derepression or heterochromatin remodeling during Met-D, indicating that the epigenetic response occurs independently of this SAM-sensing axis^31,56,58^. Likewise, pharmacologic and genetic suppression of MAT2A both failed to reproduce the effects of Met-D despite substantial loss of SAM. Together, these observations strongly suggest that methionine depletion engages a sensing mechanism that is distinct from intracellular SAM monitoring. One possibility is that methionine depletion is sensed through amino acid starvation pathways such as GCN2 activation, which detects uncharged tRNAs and drives ATF4-dependent transcriptional responses^91,92^.

Instead, Met-D was distinguished by robust activation of stress-response pathways. Only methionine deprivation induced ATF4 expression, interferon-stimulated genes, and innate immune signaling components, indicating engagement of the Integrated Stress Response. This transcriptional program coincided with derepression of multiple TE families, raising the possibility that repetitive element activation contributes to a viral mimicry-like response. Under this model, loss of heterochromatin integrity during methionine stress could permit accumulation of endogenous double-stranded nucleic acids that activate cytosolic immune sensors and amplify stress signaling. Although the present study does not establish direct causality, the close association between TE activation, innate immune signaling, and ATF4 induction suggests that chromatin remodeling and cellular stress responses are mechanistically linked during methionine deprivation.

The contribution of histone demethylases to this process appears to be secondary rather than causal. Members of the KDM4 family were transcriptionally induced specifically during Met-D, suggesting that demethylase pathways participate in the cellular response to nutrient stress. However, inhibition of KDM4 catalytic activity did not prevent H3K9me3 loss, TE induction, or ATF4 activation. Instead, KDM4 inhibition further enhanced several of these phenotypes, indicating that KDM4 activity may partially buffer chromatin instability once the stress response has been initiated. These findings argue against a simple model in which KDM4-mediated demethylation drives heterochromatin remodeling and instead implicate additional, yet unidentified mechanisms governing H3K9 methylation dynamics during nutrient limitation.

Global transcriptional profiling further reinforced the distinction between methionine and SAM depletion. Met-D induced widespread changes in gene expression involving metabolic pathways, cell cycle regulation, stress signaling, and immune responses, whereas AG-270 treatment caused minimal transcriptional disruption despite equivalent SAM depletion. The near absence of overlap between these transcriptional states indicates that Met-D creates a broader cellular adaptation that extends well beyond methyl-donor limitation. Similarly, partial SAM depletion via transient MAT2A knockdown produced heterogeneous and context-dependent transcriptional changes, likely reflecting additional non-catalytic roles of MAT2A in cellular physiology. Comparing differentially expressed genes within all conditions, there was strikingly little overlap metabolite manipulations and no significant Reactome pathways were identified. The concordance of these orthogonal approaches strengthens the conclusion that loss of methionine and loss of SAM are biologically distinct signals.

Collectively, these findings support a model in which Met-D is sensed as a broader metabolic stress that extends beyond methyl donor availability. In this framework, methionine limitation activates stress signaling pathways that become coupled to selective chromatin remodeling, leading to redistribution of H3K9 methylation, partial destabilization of heterochromatin, repetitive element expression, and large-scale transcriptional reprogramming. This response may function to preserve essential chromatin features while enabling transcriptional plasticity under nutrient stress. By contrast, isolated depletion of SAM is largely tolerated and fails to engage this adaptive program, demonstrating that methionine functions as a primary upstream regulator of epigenetic adaptation.

These insights have important implications for therapeutic strategies targeting one-carbon metabolism. Pharmacological inhibition of MAT2A, currently under clinical investigation^54^, may not recapitulate the broader epigenetic and transcriptional consequences of dietary methionine restriction. As such, the efficacy of these approaches may depend on cellular context, including methionine salvage capacity and stress-response signaling. Future studies should define how methionine depletion is sensed independently of SAM, determine the functional significance of nuclear MAT2A accumulation, and establish whether methionine-induced chromatin remodeling creates selective vulnerabilities in methionine-dependent cancers. More broadly, these findings underscore the importance of distinguishing metabolite abundance from nutrient sensing when considering how metabolism interfaces with epigenetic regulation.

## Materials and Methods

### Cell Lines

HCT116 and HEK239T cell lines were cultured at 37 °C with 5.0 % CO_2_

### Methionine Depletion of Tissue Culture Cells

HCT116 and HEK293T culture cells were originally cultured in RPMI-1640 + 10 % FBS at 37°C, 5.0 % CO_2_. The day before methionine depletion, cells were seeded at 5.0 x 10^5 for 6-well plates, or 3.5 x 10^6 for 10 cm dishes. One-hour before the depletion began, RPMI-1640 was replaced with fresh, Met-replete media. At the time of Met-depletion, cells were rinsed 2x with 3 mL of sterile DPBS pH 7.4 followed by addition of RPMI-1640 void of Met. Cells cultured for LC-MS metabolite analysis were cultured in RPMI-1640 containing 10% dialyzed FBS for these experiments.

### SAM Depletion Treatment via Drug

AG-270 was purchased through MedChemExpress® and dissolved in DMSO for a final concentration of 5 mM before aliquoting and storing at −80°C.

HCT116 and HEK293T culture cells were originally cultured in RPMI-1640 + 10 % FBS at 37°C, 5.0 % CO_2_. The day before AG-270 treatment, cells were seeded at 5.0 x 10^5 for 6-well plates, or 3.5 x 10^6 for 10 cm dishes. One-hour before SAM depletion began, RPMI-1640 was replaced with fresh, Met-replete media. At the time of inhibitor treatment, media was replaced with RPMI-1640 containing indicated concentration of AG-270.

### RNAi Mediated Inhibition

For RNAi-mediated knock-down of MAT2B and SAMTOR, 1.5 x 10^5 HCT116 cells were seeded into 6-well plates the day before transfection. The next day, a master mix of 5 µL Lipofectamine RNAiMAX was added to 120 µL Opti-MEM Reduced Serum Medium (Life Technologies, #31985070) per condition. siRNAs were diluted in a separate tube with Opti-MEM for a final working dilution of 10 mM siRNA/well. Diluted siRNAs were combined with the RNAiMAX master mix in a 1:1 ratio and incubated at room temperature for ∼15 min. 250 µL of transfection mix was added directly to wells containing previously seeded cells in 2 mL of culture medium. siMAT2B and siSAMTOR knock-down experiments were performed for 24-hours in Met-complete medium, prior to 24-hour Met depletion experiments. Knock-down efficiency was assessed via RTqPCR compared to internal house-keeping gene.

For RNAi-mediated knock-down of MAT2A, 5.0 x 10^5 HCT116 cells were reverse transfected with 30 pmol RNAi using 5 µL of Lipofectamine RNAiMAX, 500 µL Opti-MEM, and 2 mL RPMI-1640 + 10% FBS before being placed back at 37°C.

### KDM4 Inhibition

TACH101 (Zavondemstat, QC8222, TACH 101) was purchased through ProbeChem (catalog # PC-72259), resuspended in DMSO to a final concentration of 10 mM, aliquoted and stored at − 80°C.

HCT116 cells were originally cultured in RPMI-1640 + 10% FBS at 37 °C, 5.0 % CO_2_. The day before pan-KDM4 inhibition ± Met, cells were seeded at 4.0 x 10^6 in 10 cm dishes in triplicate. 40 mL RPMI ± Met was aliquoted into individual 50 mL conicals for each media change, with DMSO or drug being added directly to conicals prior to media changes. All treated media contained the same final DMSO concentration of 0.1 %. One-hour before the depletion began, RPMI-1640 was replaced with fresh RPMI-1640 media containing either DMSO, 0.5 µM, or 1 µM TACH101. At the time of Met-depletion, cells were rinsed 2x with 4 mL of sterile DPBS pH 7.4 followed by addition of RPMI-1640 void of Met. Cells were harvested after 24-hours of treatment by the following: aspirate treatment media and wash cells once with 4 mL sterile DPBS pH 7.4 before adding 2 mL Trypsin-EDTA (0.05%) and incubating at 37°C for 3-5 minutes. Once cells were lifted, trypsin was inactivated by the addition of 4 mL RPMI-1640 + 10 % FBS (with or without Met) and transferred to 15 mL conical tubes and spun 500xg for 5 minutes at 4 °C. Media was aspirated and cells were washed with 1 mL cold DPBS before transferring to 1.5 mL Eppendorf tube. 200 µL of each sample was subsequently allocated to a separate 1.5 mL Eppendorf tube for downstream RNA processing, while the larger pellet was used for histone extraction. Tubes were spun at 500xg for 5 minutes at 4°C, supernatant aspirated, and cell pellets snap frozen in liquid nitrogen before storing at −80°C.

### Histone acid-extraction

Tissue culture cell pellets were resuspended in 800 µL of ice-cold Buffer A (10 mM Tris-HCl pH 7.4, 10 mM NaCl, and 3 mM MgCl_2_) supplied with protease and histone deacetylase inhibitors (10 µg/mL leupeptin, 10 µg/mL aprotinin, 100 µM phenylmethylsulphonyl fluoride, 10 mM nicotinamide, 1 mM sodium-butyrate, and 4 µM trichostatin A) followed by 80 strokes of light pestle homogenization in a 1 mL Wheaton dounce homogenizer. Cell homogenate was then transferred to a 1.5 mL microcentrifuge Eppendorf tube and centrifugated at 800xg for 10 minutes at 4°C to pellet nuclei. The supernatant (containing cytosolic proteins) was either transferred to a fresh 1.5 mL Eppendorf tube or discarded. The nuclei pellet was resuspended in 500 mL ice-cold

DPBS pH 7.4 followed by centrifugation at 800xg for 10 minutes at 4°C. The supernatant was discarded and nuclei were again washed with 500 mL ice-cold PBS pH 7.4. Pelleted nuclei were then re-suspended in 500 mL of 0.4 N H_2_SO_4_ and rotated at 4°C for 2-hours. Samples were centrifugated at 3,400xg for 5 minutes at 4 °C to pellet nuclear debris and precipitated non-histone proteins. The supernatant was transferred to a new 1.5 mL Eppendorf tube after which 125 mL of 100% trichloroacetic acid was added and incubated overnight at 4 °C. Samples were centrifugated at 3,400xg for 5 minutes at 4°C to pellet precipitated histone proteins. The supernatant was discarded, after which the precipitant was washed with 1 mL ice-cold acetone. Samples were centrifugated at 3,400xg for 2 minutes at 4°C and the supernatant was discarded. This wash was repeated once. Residual acetone was allowed to evaporate at room temperature for 30 minutes to overnight, after which the dried precipitant was dissolved in 100 µL LC/MS-grade H_2_O. Samples were then centrifugated at 21,100xg for 5 minutes at 4°C to pellet any remaining insoluble debris. Supernatant containing purified histone was then transferred to a new 1.5 mL Eppendorf tube and stored at −20°C until future analyses. Histone PTM Analysis

To prepare purified histone proteins for LC-MS/MS analysis, 5 µg of histone were diluted with LC/MS-grade H_2_O to a final volume of 10 µL. 1 µL of 1 M triethylammonium bicarbonate was added to each sample to buffer the solution to a final pH of 7-9. Next, 1 µL of 1:50 heavy acetic anhydride:H_2_O was added to each sample and allowed to incubate for 2-minutes at room temperature. The reaction was then quenched with 1 µL of 80 mM hydroxylamine, which was allowed to incubate for 20-minutes at room temperature. Next, heavy acetylated histones were digested with 0.1 µg trypsin for 4-hours at 37 °C. Upon completion of trypsin digestion, 0.02 M NaOH was added to bring the final pH between 9-10. Anhydride-labelled trypsin-digested histone peptides were then N-terminally modified with 2 µL 4 % (v/v) phenyl-isocyanate in acetonitrile for 1-hour at 37°C. Modified peptides were desalted and eluted off of EmporeC18 extraction membrane. Eluted samples were dried completely using a ThermoFisher Scientific Savant ISS110 SpeedVac then resuspended in 30 µL sample diluent (94.9% H_2_O, 5% ACN, 0.1% TFA) and transferred to glass vials for LC-MS/MS analysis.

### RNA Extraction and cDNA Synthesis

Total RNA was extracted from HCT116 and HEK293T cells using New England Biolabs Monarch® Spin RNA Isolation Kit (Mini). Concentration and purity of RNA were determined by absorbance at 260/230 and 260/280 nm. A Monarch® Spin RNA Cleanup Kit was used to increase 260/230 ratio, if needed. A denaturing (1% bleach) 1% agarose gel was run via electrophoresis and imaged to confirm non-degraded RNA as in Aranado et. al^93^. cDNA was prepared using 0.5 - 5 ng of purified RNA and Thermo Scientific RevertAid First Strand Synthesis Kit following manufacturer’s instructions. Oligo dT primers were used in all cDNA synthesizing reactions.

### Transcript Abundance via RTqPCR

cDNA was analyzed on a Bio-Rad CFX Opus 96 Real-Time PCR System with PowerUp™ SYBR™ Green Master Mix for qPCR. 20 µL reactions were run with 2 µL diluted RNA with technical and biological replicates. Each qPCR reactions were normalized internally using primers targeting a housekeeping gene. Relative gene expression was calculated using 2^(-ΔΔCt) compared to appropriate control conditions.

### RNA Sequencing

Total RNA was extracted from HCT116 cells and assessed for purity before shipping to Novogene for RNA library synthesis and sequencing. Messenger RNA was purified from total RNA using poly-T oligo-attached magnetic beads. After fragmentation, the first strand cDNA was synthesized using random hexamer primers, followed by the second strand cDNA synthesis using either dTTP for non-strand specific library or dUTP for strand specific library. RNA libraries were prepared by Novogene followed by paired end 2×150 non-directional sequencing on their NovaSeq X Plus Series platform. All analyses were performed by Novogene using their NovoMagic interactive programming. Quality control measures indicated greater than ≥99% clean reads, of which ≥98% successfully aligned with the hg38 human genome. Replicates within groups displayed Pearson correlation coefficient (R^2^) values ≥0.99, indicating high levels of reproducibility between samples.

### Metabolite Extraction

All cells intended for LC/MS metabolite analysis were cultured in 6-well dishes with RPMI-1640 + 10 % dialyzed FBS prior to and during all treatment conditions. At time of extraction, media was aspirated and cells were quickly washed once in 1 mL ice-cold DPBS pH 7.4 before the addition of 1.5 mL of 80:20 MeOH:H_2_O extraction solvent, pre-chilled at −80°C. Plates were immediately transferred to dry ice before incubating at −80°C for 15 minutes. Next, cells were scraped off their dish and transferred to 2 mL microcentrifuge Eppendorf tube which was followed by a 5-minute, maximum speed centrifugation at 4°C. Supernatant was transferred to a new 2 mL microcentrifuge tube and dried completely using a Thermo Fisher Savant ISS110 SpeedVac. Dried metabolite extracts were resuspended in 50 µL of sample diluent (20% H_2_O, 80% ACN, 20 mM NH_4_Ac, 20 mM NH_4_OH) followed by a microcentrifugation at 4°C to pellet any insoluble debris. The supernatant was then transferred to glass vials for LC-MS analysis.

### LC-MS Metabolite Analysis

Metabolite samples resuspended in H_2_O were injected in random order onto a Thermo Fisher Scientific Vanquish UHPLC with a Waters Acquity UPLC BEH C18 column (1.7 μm, 2.1 x 100 mm; Waters Corp., Milford, MA, USA) and analyzed using a Thermo Fisher Q Exactive Orbitrap mass spectrometer in negative ionization mode. LC separation was performed over a 25-minute method with a 16-minute step-wise gradient of mobile phase (buffer A, 97% water with 3% methanol, 10 mM tributylamine, and acetic acid-adjusted pH of 8.3) and organic phase (buffer B, 100% methanol) (0 minute, 5% B; 2.5 minute, 5% B; 5 minute, 20% B; 7.5 minute, 20% B; 13 minute, 55% B; 15.5 minute, 95% B; 18.5 minute, 95% B; 19 minute, 5% B; 25 minute, 5% B, flow rate 0.2 mL/min). A quantity of 15 μL of each sample was injected into the system for analysis. The ESI settings were 30/10/1 for sheath/aux/sweep gas flow rates, 2.50 kV for spray voltage, 50 for S-lens RF level, 350°C for capillary temperature, and 300°C for auxiliary gas heater temperature. MS1 scans were operated at resolution = 70,000, scan range = 85-1250 m/z, automatic gain control target = 1 x 106, and 100 ms maximum IT.

Metabolite samples resuspended in 80 % ACN with 20 % H_2_O (20 mM final ammonium acetate and 20 mM final ammonium hydroxide) were injected in random order onto a Thermo Fisher Scientific Vanquish UHPLC with a Waters XBridge BEH Amide column (130Å, 2.5 μm, 2.1 x 150 mm; Waters Corp., Milford, MA, USA) and analyzed using a Thermo Fisher Q Exactive Orbitrap mass spectrometer in positive ionization mode. LC separation was performed over a 22-minute method with a 13-minute step-wise gradient of mobile phase (buffer A, 5% ACN with 95% methanol, 20 mM ammonium acetate and 20 mM ammonium hydroxide) and organic phase (buffer B, 80% ACN with 20% H_2_O, 20 mM ammonium acetate and 20 mM ammonium hydroxide) (0 minute, 100% B; 3 minute, 100% B; 3.2 minute, 90% B; 6.2 minute, 90% B; 6.5 minute, 80% B; 10.5 minute, 80% B; 10.7 minute, 70% B; 13.5 minute, 70% B; 13.7 minute, 45% B; 16 minute, 45% B; 16.5 minute, 100% B; 22 minute, 100% B; flow rate 0.3 mL/min). A quantity of 15 μL of each sample was injected into the system for analysis. The ESI settings were 10/5/1 for sheath/aux/sweep gas flow rates, 3.50 kV for spray voltage, 50 for S-lens RF level, 350°C for capillary temperature, and 30°C for auxiliary gas heater temperature. MS1 scans were operated at resolution = 70,000, scan range = 60-186 m/z and 187-900 m/z, automatic gain control target = 3 x 106, and 200 ms maximum IT.

Raw data files were converted into mzml for metabolite identification and peak AreaTop quantification using El-MAVEN (v0.12.1-beta)

### Western blotting

Proteins separated on SDS-PAGE gels were transferred onto 0.2 mm nitrocellulose membrane using a wet transfer method, except in the case of histone proteins which were transferred via a Bio-Rad Trans-Blot Turbo Transfer System using the Standard SD protocol. Membranes were ponceau stained to ensure even transfer before incubating with 5-10 mL Intercept Blocking Buffer (TBS) from LICORbio for 1-hour at RT. Primary antibodies were diluted in 1:1 mix of TBS-T : Intercept Blocking Buffer (TBS) and incubated with membranes overnight at 4°C. Membranes were washed 3x in fresh TBST for 10-minutes each. LICOR secondary antibodies were used at 1:15,000 – 1:30,000 dilution in 1:1 TBS-T : Intercept Blocking Buffer for 1-hour incubation at RT. Membranes were washed 3x in fresh TBS-T for 10-minutes each, followed by 1x wash in TBS.

Membranes were imaged using an Odyssey Infrared Imager (model no. 9120). Densitometry was performed using Image Studio Lite software. All data was normalized to an internal loading control.

### MAT2A and MAT2B Recombinant Protein Purification

The *H. sapiens* MAT2A (MAT2AA-c001, Addgene plasmid # 53648) *E. coli* expression plasmid used in this study was acquired from Addgene, made possible by a gift from Nicola Burgess-Brown. The *H. sapiens* MAT2B gBlock was cloned into a pEQEOL backbone using BamHI and SalI restriction enzymes.

Transformed Rosetta™ E. coli competent cells were cultured in 1 L of 2xYT media at 37 °C to an O.D. 600 of 0.6 - 0.8. IPTG was then added to each culture at a final concentration of 0.5 mM. Cultures were allowed to grow for 16-hours at 18°C before harvested and stored at −80°C. Pellets were resuspended in 30 mL Buffer A (50 mM NaPi pH 7.2, 250 mM NaCl, 10 mM imidazole, 1 mM 2-Mercaptoethanol) and sonicated on ice in the presence of lysozyme, DNAse I, and protease inhibitors (aprotinin, leupeptin, pepstatin A, PMSF). Lysate was centrifuged at 18,000 rpm for 45 min, and the supernatant was collected and passed through a 0.8 µm SCFA syringe filter.

Filtered supernatant was then loaded onto a HisTrap FF nickel column in line with a GE AKTA FPLC using Buffer A. Non-specific proteins were flushed from the column using a 0-25 % gradient in Buffer B (50 mM NaPi pH 7.2, 250 mM NaCl, 250 mM imidazole, 1 mM 2-Mercaptoethanol) for 10 min, followed by a 25% Buffer B hold for ∼30 min. Remaining proteins were eluted using a 25-100 % gradient in Buffer B for 40 min and collected in time-dependent fractions. Fractions were analyzed by Coomassie staining of SDS-PAGE gels to assess purity.

Fractions determined to contain protein of interest were pooled, concentrated (Millipore Sigma AMICON ULTRA-15), and dialyzed overnight at 4°C in 4 L of dialysis buffer (20 mM HEPES pH 7.5, 150 mM KCl, 10 mM MgCl_2_ hexahydrate, 1 mM TCEP, 10 % glycerol). Following dialysis, precipitated protein was pelleted via centrifugation at 4°C for 10 min at 8,000xg and supernatant was collected. Protein concentration determined via A280 absorbance values and extinction coefficient correction. Final protein samples were aliquoted into single-use 0.2 mL PCR tubes and flash frozen in liquid nitrogen prior to long-term storage at −80°C.

### Fluorescent Polarization

Recombinant proteins were thawed on ice and diluted to a concentration of 200 µM in dialysis buffer (20 mM HEPES pH 7.5, 150 mM KCl, 10 mM MgCl_2_ hexahydrate, 1 mM TCEP, 10 % glycerol) before serially diluting and aliquoting 15 µL of protein into 384-well black flat bottom non-binding plates. Next, 5 µL of 100 nM FAM-labelled Widom DNA (10-, 20-, or 30-basepair fragments) was added to reactions in triplicate and mixed well by pipetting. The plate was protected from light and centrifuged briefly before fluorescence was read on a PheraStar plate reader.

### Electromobility Shift Assay (EMSA)

10 mL non-denaturing gels were prepared in-house and poured into siliconized glass plates for polymerization ahead of time. Recombinant proteins were serially diluted, mixed as 10 µL reactions, and allowed to incubate for 10 min at RT. Reaction mixes were loaded onto native gels and separated via electrophoresis on ice. Gels were stained with SYBR safe DNA dye and imaged on a Typhoon Imager.

### Subcellular Fractionation Methods

*Dounce homogenation*: Tissue culture cell pellets were resuspended in 400 µL of ice-cold Buffer A (10 mM Tris-HCl pH 7.4, 10 mM NaCl, and 3 mM MgCl_2_) supplied with protease and phosphatase inhibitor tablets (Thermo Scientific A32961) followed by 80 strokes of light pestle homogenization in a 1 mL Wheaton dounce homogenizer. Cell homogenate was then transferred to a 1.5 mL microcentrifuge Eppendorf tube. The used dounce homogenizer was rinsed with an additional 100 µL which was then combined with cell homogenate and centrifugated at 800xg for 10-minutes at 4°C to pellet nuclei. Supernatant (cytoplasm) was removed and transferred to a new 1.5 mL microcentrifuge tube and centrifugated at 2,000xg for 10-minutes at 4°C to pellet insoluble material, before transferring supernatant to a fresh 1.5 mL microcentrifuge tube and stored at − 80°C. The nuclear pellets were washed twice in 500 µL ice-cold PBS and spun at 800xg for 5-minutes at 4°C. Nuclei were next solubilized in 200 µL RIPA buffer (150 mM NaCl, 1.0%(v/v) NP-40, 0.5 % (w/v) Na-deoxycholate, 0.1 % (v/v) SDS, and 50 mM Tris pH 8.0) with protease and phosphatase inhibitors and left to incubate on ice for 30 minutes, vortexing nuclei every 10 min. Solubilized nuclei were centrifugated at 14,000xg for 20 minutes and transferred to a fresh 1.5 mL microcentrifuge tube before long-term storage at −80°C.

*NE-PER Kit*: A Thermo Scientific NE-PER Nuclear and Cytoplasmic Extraction Kit (#78833) was used according to manufacturer recommendations on previously flash-frozen cell pellets.

*Lyse-and-Wash*: A modified “lyse-and-wash” subcellular fractionation protocol was adapted from Senichkin et al. (2021). Flash-frozen cell pellets were resuspended in freshly-made ice-cold hypotonic buffer (20 mM Tris-HCl pH 7.4, 10 mM KCl, 2 mM MgCl_2_, 1 mM EGTA, 0.5 mM dithiothreitol, 0.5 mM AEBSF) and incubated on ice for 3 min. NP-40 was added to resuspended cells at a final concentration of 0.1%, mixed gently, and incubated on ice for 3-minutes before centrifugation at 1,000xg for 5-minutes at 4°C to separate nuclei (pellet) from cytosol (supernatant). Supernatant was transferred to a new 1.5 mL microcentrifuge tube and placed on ice prior to high-speed centrifugation (15,000xg) for 3-minutes at 4°C to pellet insoluble protein. Nuclei pellets were resuspended by pipetting in isotonic buffer (20 mM Tris-HCl pH 7.4, 150 mM KCl, 2 mM MgCl_2_, 1 mM EGTA, 0.5 mM dithiothreitol, 0.5 mM AEBSF, 0.1 % NP-40) and incubated on ice for 5-10 minutes prior to centrifugation at 1,000xg for 3-minutes at 4°C. To separate soluble versus insoluble nuclear proteins, nuclei were incubated on ice for 20-30 minutes in ice-cold RIPA buffer (25 mM Tris-HCl pH 7.4, 150 mM NaCl, 0.1 % SDS, 0.5 % Na-deoxycholate, 1.0 % NP-40) supplemented with protease and phosphatase inhibitors, with vortexing every 10-minutes. Next, nuclei were centrifugated at 2,000xg for 3-minutes at 4°C to separate RIPA-soluble (supernatant) versus RIPA-insoluble (pellet) nuclear proteins. RIPA-soluble supernatant was transferred to a new 1.5 mL microcentrifuge tube, while the insoluble pellet was incubated with DNAse I to degrade chromatin and free remaining proteins.

### Serum Starvation

3.5 x 10^5 cells were seeded into 10 cm plates 1-day prior to serum starvation. One-hour prior to starvation, fresh medium was given and cells were placed back at 37°C. Next, plates were rinsed twice in DPBS before RPMI-1640 without FBS addition, followed by 24-hours of growth in serum starved media. Cells were harvested after starvation, or medium was changed to fresh RPMI-1640 + 10% FBS before harvesting at 1- and 3-hours post supplementation.

## Supporting Information (attached separately)

Table S1: Antibodies for Western Blotting

Table S2: RTqPCR Oligos

Table S3: siRNA Transfection Materials

Table S4: Reagents

Table S5: Differentially Expressed Genes

Table S6: Reactome Analysis of Shared DEGs

## Acknowledgments

The authors would like to acknowledge the use of NovoMagic, a platform by Novogene, for all RNA-seq plots and analysis. Models created with BioRender.

## Author Contributions

CML conceptualized project, planned and conducted experiments, analyzed results, and wrote the manuscript. SAH aided with technical and statistical approaches. CML, SAH, and JMD contributed to data interpretation and manuscript edits.

