## Supplemental Information for "Methionine, Not *S*-adenosylmethionine, Acts as a Primary Metabolic Stress Signal for Chromatin Remodeling"

(Supplementary Information) Figure S1 - MAT2A inhibition does not phenocopy heterochromatin adaptation in a conserved manner.

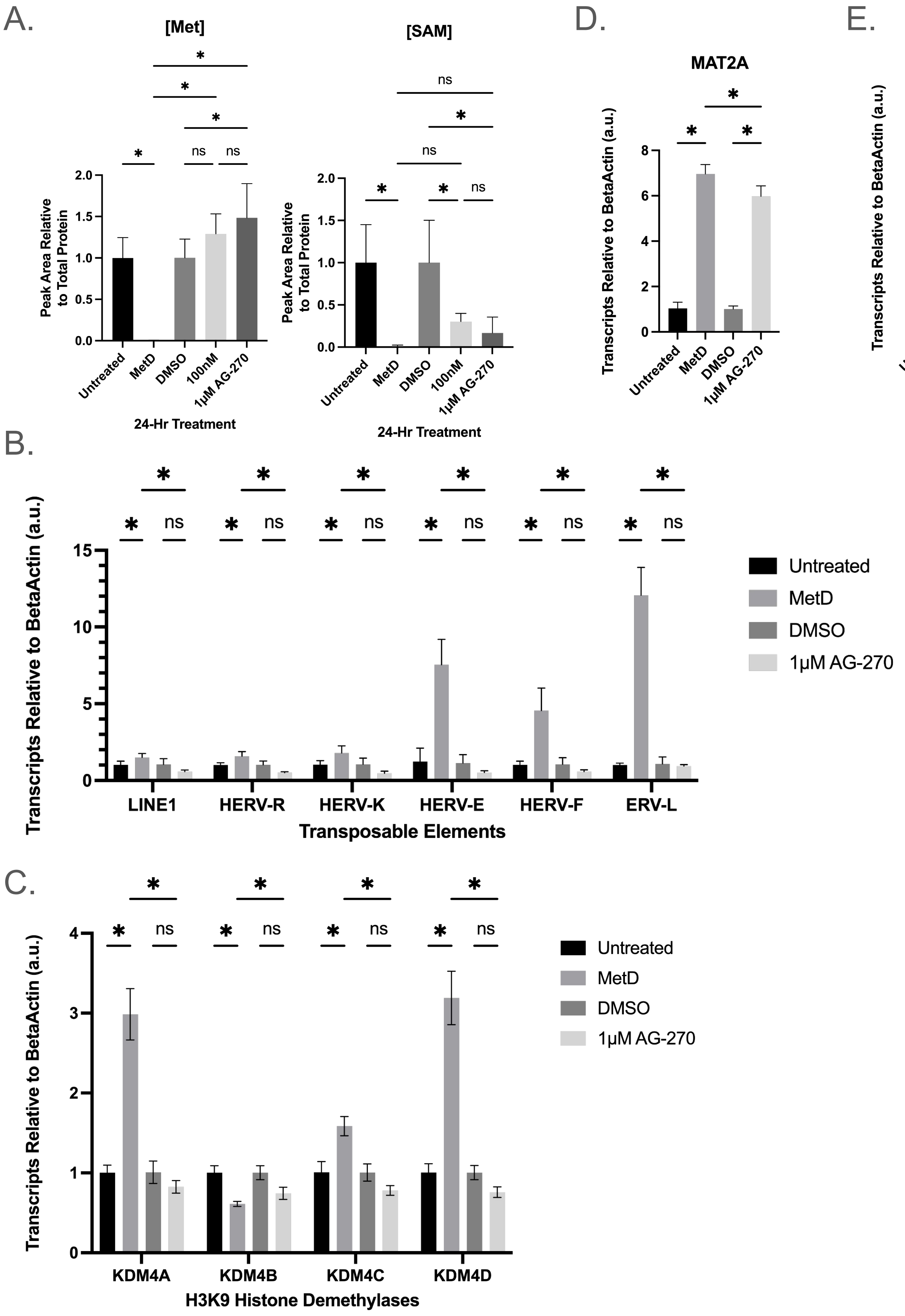

**(Supplementary Information) Figure S2 - SAMTOR, the SAM-sensing negative regulator of mTORC1, is dispensable for methionine-depletion mediated epigenetic adaptation.**

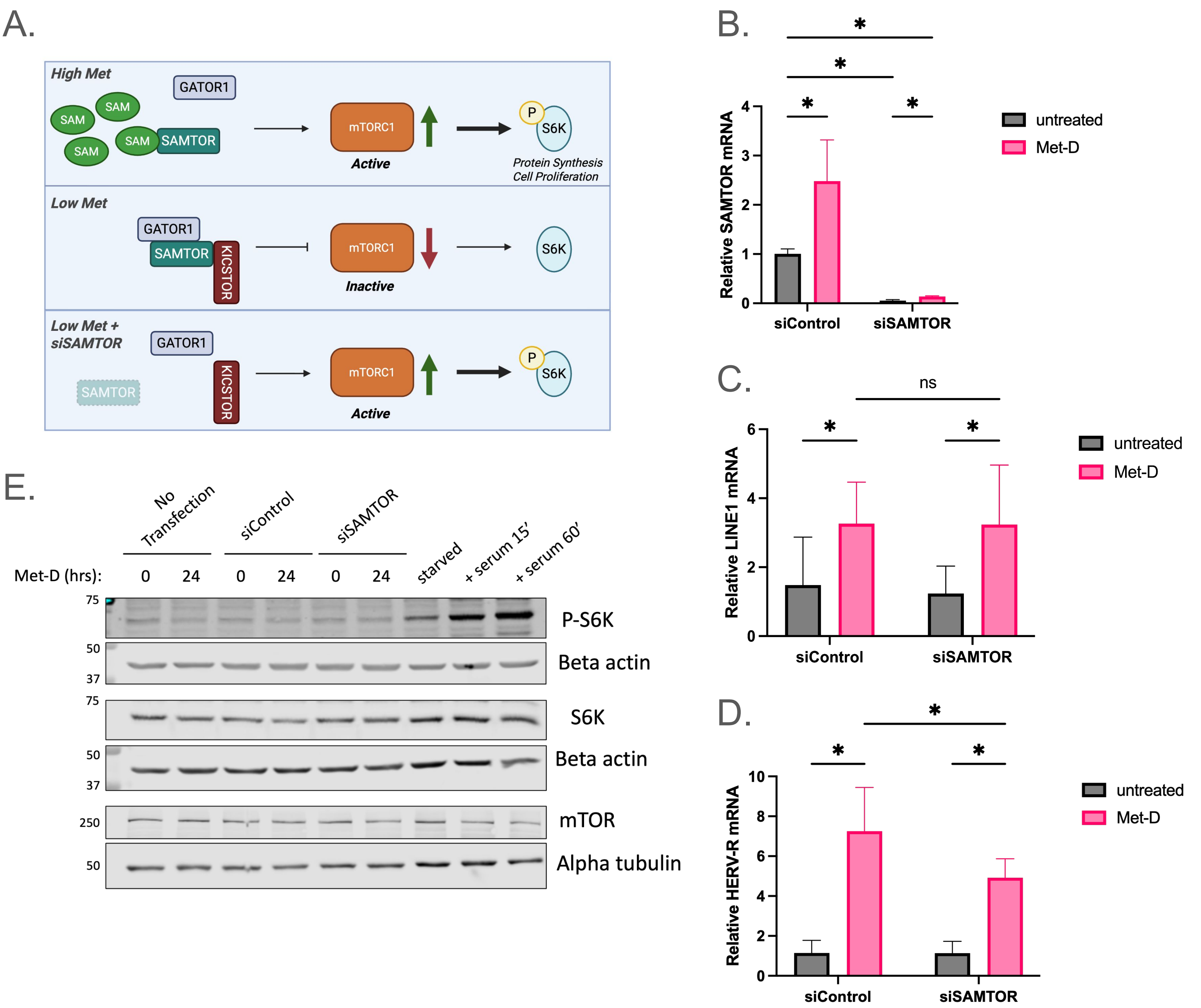

**(Supporting Information) Figure S3 - Methionine depletion activates an innate immune response consistent with viral mimicry.**

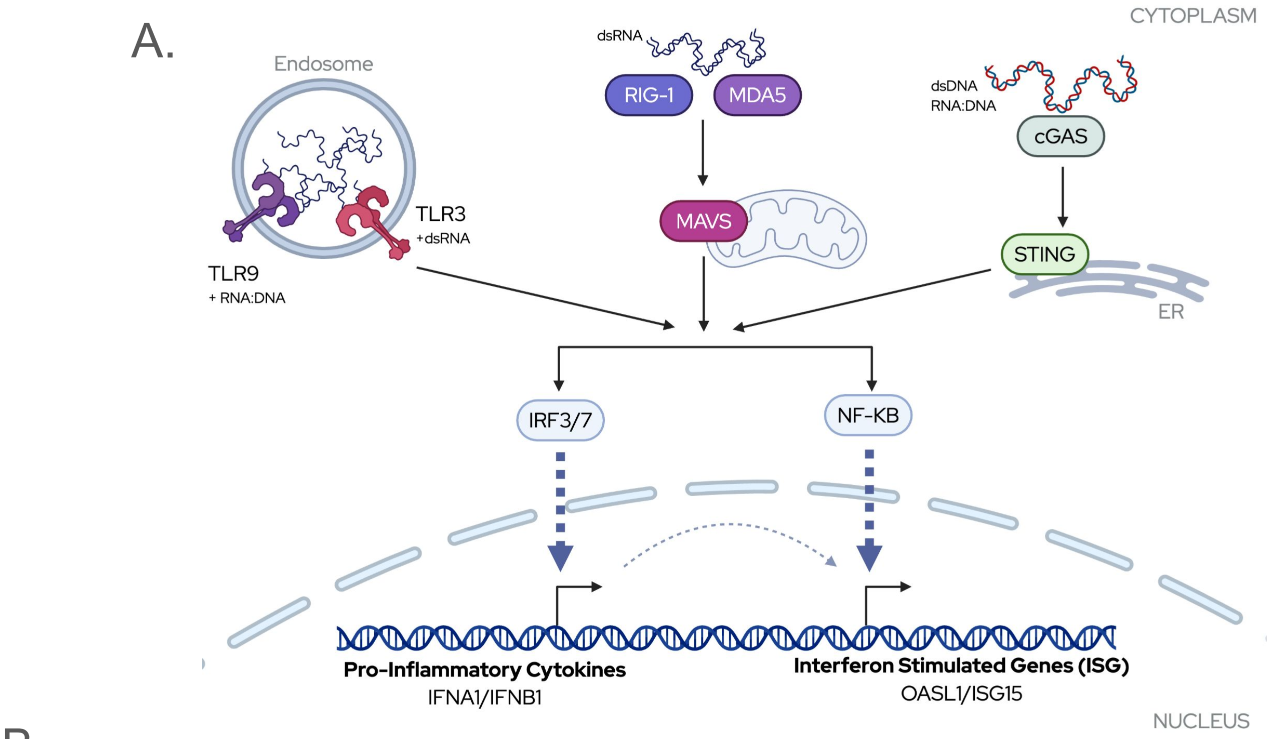


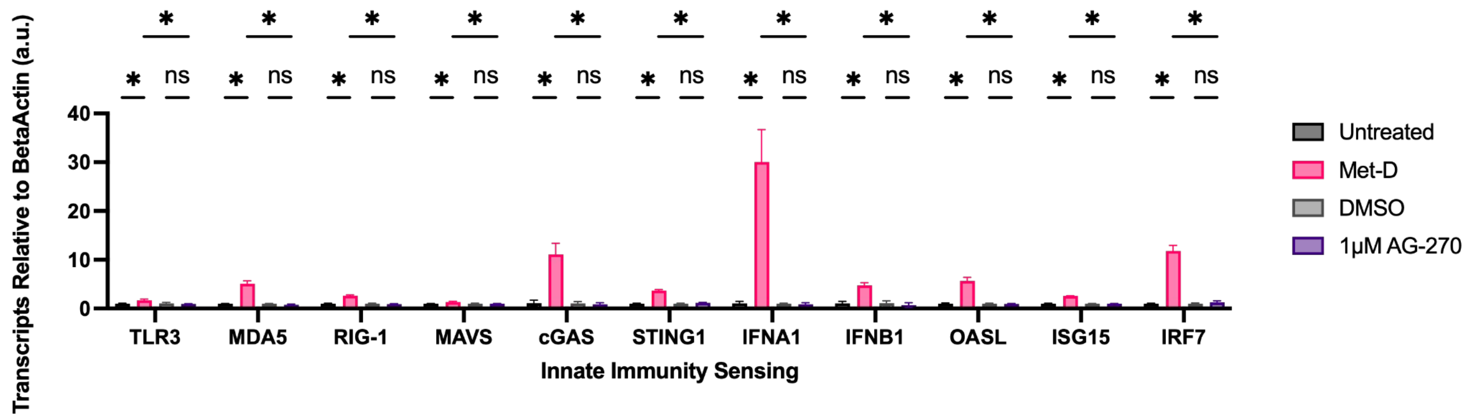

### (Supporting Information) Figure S4 - MAT2A knock-down reduces SAM but produces a mixed heterochromatin phenotype and distinct transcriptional outcomes.

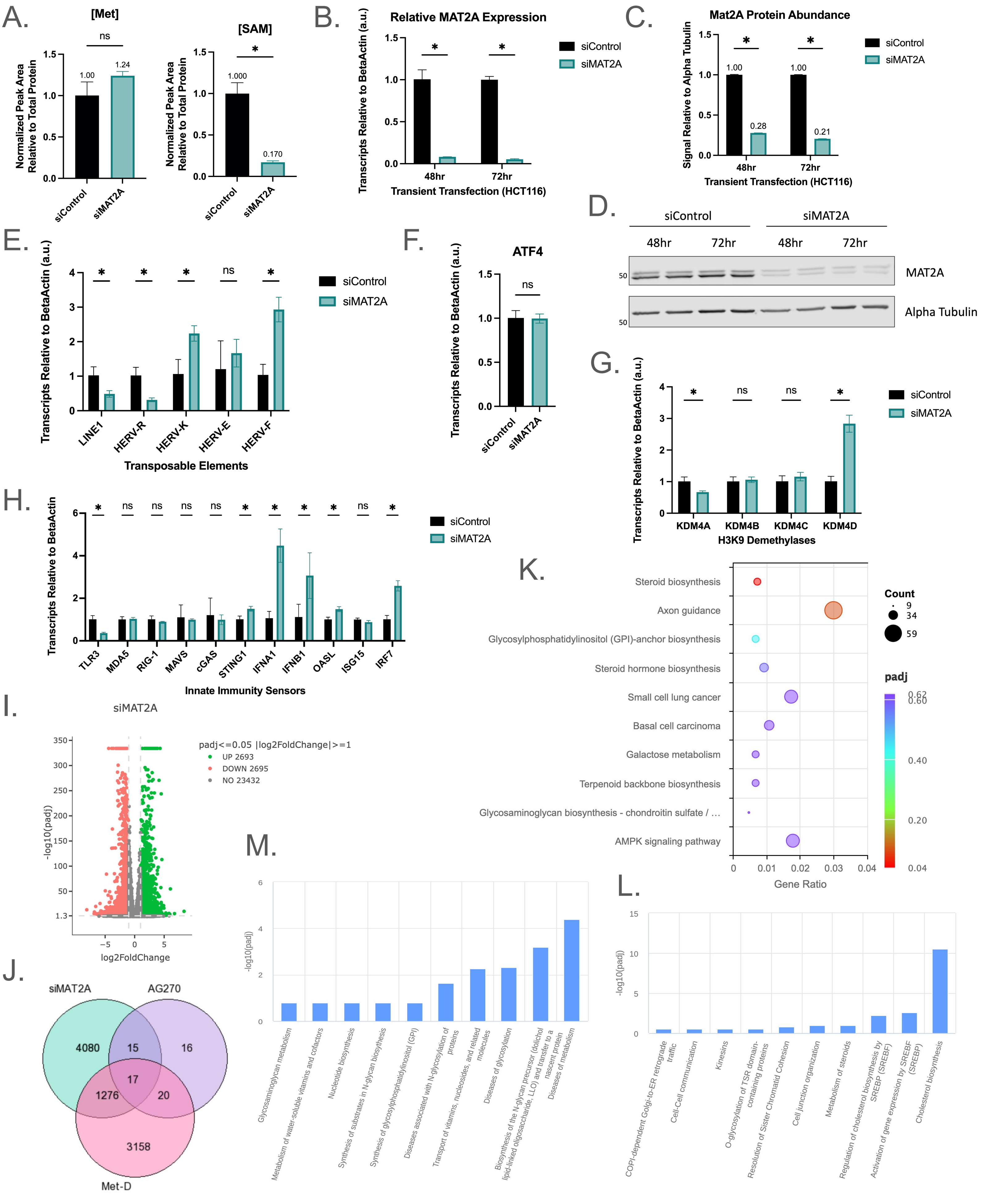

### Supporting Information

**Figure S1: MAT2A inhibition does not phenocopy heterochromatin adaptation in a conserved manner.**

(**A**) HEK293T metabolite abundance of relative intracellular Met and SAM compared to appropriate control condition following 24-hours of growth in Met-D or 1µM AG-270 treated medium. n=3; error bars represent SD; $*p < 0.05$ (Ordinary one-way ANOVA). (**B-E**) Bar graphs depicting HEK293T transcript abundance of indicated genes relative to internal housekeeping and normalized to the appropriate DMSO-matched control following 24-hours of indicated treatment. n=3 biological replicates; error bars represent SD; $*p < 0.05$ (2way ANOVA).

**Figure S2: SAMTOR, the SAM-sensing negative regulator of mTORC1, is dispensable for methionine-depletion mediated epigenetic adaptation.**

(**A**) Model of SAMTOR activation based on published literature[57,95]. Under high Met/SAM conditions, SAMTOR binds SAM ($K_d$ <10 µM), inducing a conformational change which prevents its association with, and inhibition of, mTORC1 (mechanistic target of rapamycin complex 1). S6K is a canonical downstream target of mTORC1 and is phosphorylated when the complex is active. When intracellular SAM concentrations drop below the kinetic threshold conducive for binding, unbound SAMTOR associates with negative regulators of mTORC1 (i.e. GATOR1/KICSTOR) leading to inhibition of mTORC1, unphosphorylated S6K, and the blockage of growth signals. By knocking-down SAMTOR under methionine limited conditions, this theoretically removes the cellular "brake" leading to constitutive mTORC1 activation (and S6K phosphorylation) under nutrient stress. (**B**) RTqPCR analysis of SAMTOR transcripts relative to internal housekeeping gene, following 48-hours of siSAMTOR and 24-hours of Met-D. n=3; error bars represent SD; *p value < 0.05 (2way ANOVA). Note that there were no SAMTOR-specific antibodies available at the time of this study to assess SAMTOR protein levels. (**C-D**) RTqPCR analysis of TE expression levels following 48-hours siSAMTOR and 24-hours of Met-D relative to internal housekeeping gene. n=3; error bars represent SD; *p value < 0.05 (2way ANOVA). (**E**) Western blot analysis of HCT116 cells subjected to 48-hours of siSAMTOR and 24-hours Met-D (left). Phosphorylated-S6K (P-S6K) was used as a read-out for mTORC1 activation. As a positive control for P-S6K, HCT116 cells were serum-starved for 24-hours prior to harvest, or given serum-containing medium for 15 or 60 min before harvesting. Beta Actin and Alpha Tubulin used as loading controls, molecular weight markers indicated on left. Representative image, n=2.

**Figure S3: Methionine depletion activates an innate immune response consistent with viral mimicry.**

(**A**) Model of activation of the innate immunity sensors, which are triggered by double-stranded RNA (dsRNA), RNA:DNA hybrids, or viral dsDNA[68,96,97]. Cytoplasmic nucleotide sensing proteins (TLR3, RIG-1, MDA5, MAVS, cGAS, STING1) converge to activate transcription factors such as IRF3/7 and NF-KB for nuclear translocation and expression of Pro-Inflammatory Cytokines (IFNA1 and IFNA2) and Interferon Stimulated Genes (ISGs; OASL and ISG15). (**B**) Bar graph depicting transcript abundance of innate immune-sensing proteins relative to internal housekeeping genes and normalized to the appropriate DMSO-matched control following 24-hours of indicated treatment. n=3 biological replicates; error bars represent SD; *p < 0.05 (2way ANOVA).

**Figure S4: MAT2A knock-down reduces SAM, but produces a mixed heterochromatin phenotype and distinct transcriptional outcomes.**

(**A**) Metabolite abundance of intracellular SAM and Met relative to non-targeting siRNA controls following 48-hours of siMAT2A knock-down. n=3; error bars represent SD; $*p < 0.05$ (Ordinary one-way ANOVA). (**B**) RTqPCR analysis of MAT2A transcript abundance relative to internal housekeeping gene, following 24- or 48-hour siRNA knock-down in HCT116 cells. n=3; error bars represent SD; $*p$ value $< 0.05$ (Welch's t-Test). (**C**) Bar graph of quantified MAT2A protein abundance relative to loading, following 48- or 72-hours of transient knock-down depicted in (D). n=2; error bars indicate SD. (**D**) Representative Western blot of whole-cell MAT2A levels following 48- or 72-hour knock-down. Alpha Tubulin used as loading control. (**E-H**) RTqPCR analysis of indicated transcripts relative to internal housekeeping gene, normalized to non-targeting siRNA control. n≥3; error bars represent SD; $*p$ value $< 0.05$ (Welch's t-Test). (**I**) Volcano plot of significant and differentially expressed genes following RNA sequencing of transient MAT2A knock-down compared to non-targeting siRNA control. The x-axis represents fold-change in gene expression between samples, and y-axis indicates the statistical significance of the differences. Each dot represents a gene, while red indicates significant down-regulation and green indicates significant up- regulation. padj ≤ 0.05; log2 fold-change cutoff ≥ 1. (**J**) Venn diagram depicting shared and unique differentially regulated genes between indicated treatments. Specific gene lists can be found in Supporting Information. (**K**) KEGG enrichment analysis of pathways altered by MAT2A knock-down. Bubble colors indicate p-adj and bubble size indicates the number of differentially expressed genes identified within the KEGG pathway. Steroid biosynthesis is the only significantly altered KEGG pathway. (**L**) Reactome pathway analysis of down-regulated processes and (**M**) up-regulated processes following 48-hours of siMAT2A knock-down. For pathway enrichment, Reactome uses a threshold of $padj < 0.05$ to determine significant enrichment.

**Table S1 - Antibodies for Western Blotting**

| PRIMARY | DILUTION | PURCHASE SOURCE | CATALOG # |
|---|---|---|---|
| **Alpha Tubulin (11H10)** | 1:1000 | Cell Signaling Technology | 2125S |
| **Alpha Tubulin** | 1:1000 | Cell Signaling Technology | 3873S |
| **Alpha Tubulin** | 1:1000 - 1:3000 | Abcam | ab176560 |
| **Beta Actin (AC-15)** | 1:1000 | Thermo Fisher Scientific | AM4302 |
| **Histone H3 (96C10)** | 1:1000 - 1:3000 | Cell Signaling Technology | 3638S |
| **Histone H3** | 1:1000 | Abcam | ab76307 |
| **H3K9ac** | 1:1000 | Millipore | 07-352 |
| **H3K9me0** | 1:1000 | Active Motif | 91155 |
| **H3K9me1** | 1:1000 | Invitrogen | MA5-33385 |
| **H3K9me3 (D4W1U)** | 1:1000 | Cell Signaling Technology | 13969T |
| **Histone H3** | 1:1000 | Abcam | ab76307 |
| **Histone H3 (96C10)** | 1:1000 - 1:3000 | Cell Signaling Technology | 3638S |
| **Lamin B1 (A-11)** | 1:1000 | Santa Cruz Biotechnology | sc-377000 |
| **Lamin B1 (D9V6H)** | 1:1000 | Cell Signaling Technology | 13435S |
| **MAT2A (3A2) - BSA Free** | 1:1000 | Novus Biologicals | NBP1-28605 |
| **MAT2B (16H3L1)** | 1:500 - 1:1000 | Life Technologies | 703221 |
| **mTOR (7C10)** | 1:1000 | Cell Signaling Technology | S983S |
| **P70 S6K (49D7)** | 1:1000 | Cell Signaling Technology | 2708S |
| **Phospho-p70 S6K (Thr389)** | 1:1000 | Cell Signaling Technology | 9234S |
| SECONDARY | DILUTION | PURCHASE SOURCE | CATALOG # |
| **IRDye® 680RD Goat anti-Mouse IgG** | 1:15,000 - 1:30,000 | LICOR Bio | 925-68070 |
| **IRDye® 680RD Goat anti-Rabbit IgG** | 1:15,000 - 1:30,000 | LICOR Bio | 926-68071 |
| **IRDye® 800CW Anti-Mouse IgG** | 1:15,000 - 1:30,000 | LICOR Bio | 925-32210 |
| **IRDye® 800CW Anti-Rabbit IgG** | 1:15,000 - 1:30,000 | LICOR Bio | 926-32211 |

**Table S2 - RTqPCR Oligo Sequences (*Homo sapiens*)**
*** indicates house-keeping gene used for normalization**

| Gene Target | Forward Primer (3' → 5') | Reverse Primer (3' → 5') | Source |
|---|---|---|---|
| **Beta Actin *** | CATGTACGTTGCTATCCAGGC | CTCCTTAATGTCACGCACGAT | Zhao, LM. et al (Asian Pac J Cancer Prev, 2014) |
| **GAPDH *** | TTGGCTACAGCAACAGGGTG | GGGGAGATTCAGTGTGGTGG | Kwon, N. et al (Gene Reports, 2021) |
| **LINE1** | ACAGCTTTGAAGAGAGCAGTGGTT | AGTCTGCCCGTTCTCAGATCT | Haws, S.A. et al (Cell Metab, 2020) |
| **HERV-R** | CATGGGAAGCAAGGGAACT | CTTTCCCCAGCGAGCAATAC | Haws, S.A. et al (Cell Metab, 2020) |
| **HERV-K** | GGCCATCAGAGTCTAAACCACG | CTGACTTTCTGGGGGTGGCCG | Haws, S.A. et al (Cell Metab, 2020) |
| **HERV-E** | GGTGTCACTACTCAATACAC | GCAGCCTAGGTCTCTGG | Shen, J.Z. et al (Cell, 2021) |
| **HERV-F** | CCTCCAGTCACAACAACTC | TATTGAAGAAGGCGGCTGG | Shen, J.Z. et al (Cell, 2021) |
| **ERV-L** | ATATCCTGCCTGGATGGGGT | GAGCTTCTTAGTCCTCCTGTGT | Shen, J.Z. et al (Cell, 2021) |
| **MAT2A** | CCACGAGGCGTTCATCGAGG | AAGTCTTGTAGTCAAAACCT | Shiraki, N. et al (Cell Metab, 2014) |
| **MAT2B** | TGGGGAGCACTTGAAAGAG | CTTAGCGGCAACATGGG | Shiraki, N. et al (Cell Metab, 2014) |
| **SAMTOR** | CCAAAATGGTGGGAAGAGAA | GCAGCTGCCAACATCAAGTA | Gift from Vince Cyns lab (UW-Madison) |
| **ATF4** | CCCTTCACCTTCTTACAACCTC | TGCCCAGCTCTAAACTAAAGGA | Stone, K. et al (Scientific Reports, 2021) |
| **FGF21** | GGGAGTCAAGACATCCAGGT | GGCTTCGGACTGGTAAACAT | Stone, K. et al (Scientific Reports, 2021) |
| **KDM4A** | TGCGGCAAGTTGAGGATGGTCT | GCTGCTTGTTCTTCCTCCTCATC | Zhao, E. et al (Cell Press, 2016) |
| **KDM4B** | GCCGAGAGGAAGTTCAACGCAG | TGCCTCCTTCTCAGTCTGTAGG | Zhao, E. et al (Cell Press, 2016) |
| **KDM4C** | CCGATGACTCTTGTGAAGCAGC | GACTTCGTCTGCCAAAGGTGGA | Zhao, E. et al (Cell Press, 2016) |
| **KDM4D** | CCTGAACGCTATGACCTGTGGA | TCTCCTGGGTAACTGGACTTCC | Zhao, E. et al (Cell Press, 2016) |
| **TLR3** | TGGTTGGGCCACCTAGAAGTA | TCTCCATTCCTGGCCTGTG | Zhang, S.M. et al (Nature, 2021) |
| **MDA5 (IFIH1)** | GAGCAACTTCTTTCAACCACAG | CACTTCCTTCTGCCAAACTTG | Zhang, S.M. et al (Nature, 2021) |
| **RIG-1 (DDX58)** | CCAGCATTACTAGTCAGAAGGAA | CACAGTGCAATCTTGTCATCC | Zhang, S.M. et al (Nature, 2021) |
| **MAVS** | AGGAGACAGATGGAGACACA | CAGAACTGGGCAGTACCC | Zhang, S.M. et al (Nature, 2021) |
| **cGAS** | TAACCCTGGCTTTGGAATCAAAA | TGGGTACAAGGTAAAATGGCTTT | Zhang, S.M. et al (Nature, 2021) |
| **STING1** | AGCATTACAACAACCTGCTACG | GTTGGGGTCAGCCATACTCAG | Zhang, S.M. et al (Nature, 2021) |
| **IFNA1** | AATGACAGAATTCATGAAAGCGT | GGAGGTTGTCAGAGCAGA | Zhang, S.M. et al (Nature, 2021) |
| **IFNB1** | GCCATCAGTCACTTAAACAGC | GAAACTGAAGATCTCCTAGCCT | Zhang, S.M. et al (Nature, 2021) |
| **OASL** | GCAGAAATTTCCAGGACCAC | CCCATCACGGTCACCATTG | Zhang, S.M. et al |

| | | | |
|---|---|---|---|
| | | | (Nature, 2021) |
| **ISG15** | CCTTCAGCTCTGACACC | CGAACTCATCTTTGCCAGTACA | Zhang, S.M. et al (Nature, 2021) |
| **IRF7** | GTGGACTGAGGGCTTGTAG | TCAACACCTGTGACTTCATGT | Zhang, S.M. et al (Nature, 2021) |
| **FoxM1** | ACTTTAAGCACATTGCCAAGC | CGTGCAGGGAAAGGTTGT | Zhao, E. et al (Cell Press, 2016) |
| **cMyc** | AAACACAAACTTGAACAGCTAC | ATTTGAGGCAGTTTACATTATGG | Villa, E. et al (MolCell, 2021) |

**Table S3 - siRNA Transfection Reagents**

| GENE | siRNA TYPE | SOURCE |
|---|---|---|
| Human MAT2A | Stealth RNAi Sense | Thermo Fisher (Kera et al., 2013) |
| Human Control siRNA | Stealth RNAi Sense | Thermo Fisher (Kera et al., 2013) |
| Human MAT2B | ON-TARGET plus SMARTpool siRNA (27430) | Horizon Dharmacon |
| Human BMT2 (aka SAMTOR) | ON-TARGET plus SMARTpool siRNA (154743) | Horizon Dharmacon |
| Human Non-Targeting | siGENOME - Pool #1 | Horizon Dharmacon |

**Table S4 – Reagents**

| REAGENT | SOURCE | CATALOG # |
|---|---|---|
| AG-270 | MedChem Express | HY-138630 |
| Dialyzed Fetal Bovine Serum (FBS) | Fisher Scientific | SH3007903 |
| Dimethyl Sulfoxide (DMSO) | Santa Cruz Biotechnology | sc-358801 |
| DPBS, no calcium, no magnesium | Thermo Fisher Scientific | 14190250 |
| Fetal Bovine Serum (FBS), Premium | Thermo Scientific | A5670701 |
| HCT116 *H. sapiens* cell line | ATCC | CCL-247 |
| HEK293T *H. sapiens* cell line | ATCC | CRL-3216 |
| Trypsin-EDTA (0.05%), phenol | Thermo Scientific | 25300062 |
| Intercept® (TBS) Blocking Buffer | LICORbio | 927-60001 |
| Lipofectamine RNAiMAX | Thermo Scientific | 13778150 |
| Monarch® Spin RNA Cleanup Kit | New England Biolabs | T2030L |
| Monarch® Spin RNA Isolation Kit (Mini) | New England Biolabs | T2110S |
| NE-PER Nuclear and Cytoplasmic Extraction Kit | Thermo Scientific | 78833 |
| Opti-MEM Reduced Serum Medium | Life Technologies | 31985070 |
| PowerUp™ SYBR™ Green Master Mix for qPCR | Applied Biosciences | A25742 |
| RevertAid First Strand Synthesis Kit | Thermo Scientific | K1622 |
| Rosetta ™ 2 Competent Cells | Sigma Corporation of America | 71402-3 |
| RPMI-1640 Media | Thermo Scientific | 11875119 |
| RPMI no methionine Media | Life Technologies | A1451701 |
| TACH101 (Zavondemstat) | ProbeChem | PC-72259 |
| Trypsin/Lys-C Mix, Mass Spec Grade | Promega | V5071 |
